# Ensemble learning and ground-truth validation of synaptic connectivity inferred from spike trains

**DOI:** 10.1101/2024.02.01.578336

**Authors:** Christian Donner, Julian Bartram, Philipp Hornauer, Taehoon Kim, Damian Roqueiro, Andreas Hierlemann, Guillaume Obozinski, Manuel Schröter

## Abstract

Probing the architecture of neuronal circuits and the principles that underlie their functional organization remains an important challenge of modern neurosciences. This holds true, in particular, for the inference of neuronal connectivity from large-scale extracellular recordings. Despite the popularity of this approach and a number of elaborate methods to reconstruct networks, the degree to which synaptic connections can be reconstructed from spike-train recordings alone remains controversial. Here, we provide a framework to probe and compare connectivity inference algorithms, using a combination of synthetic and empirical ground-truth data sets, obtained from simulations and parallel single-cell patch-clamp and high-density microelectrode array (HD-MEA) recordings in vitro. We find that reconstruction performance critically depends on the regularity of the recorded spontaneous activity, i.e., their dynamical regime, the type of connectivity, and the amount of available spike train data. We find gross differences between different algorithms, and many algorithms have difficulties in detecting inhibitory connections. We therefore introduce an ensemble artificial neural network (eANN) to improve connectivity inference. We train the eANN on the validated outputs of six established inference algorithms, and show how it improves network reconstruction accuracy and robustness. Overall, the eANN was robust across different dynamical regimes, with shorter recording time, and ameliorated the identification of synaptic connections, in particular inhibitory ones. Results indicated that the eANN also improved the topological characterization of neuronal networks. The presented methodology contributes to advancing the performance of inference algorithms and facilitates our understanding of how neuronal activity relates to synaptic connectivity.

**Author summary:** This study introduces an ensemble artificial neural network (eANN) to infer neuronal connectivity from multi-unit spike time recordings. We compare the eANN to previous algorithms and validate it using simulations and HD-MEA/patch-clamp datasets. The latter is obtained from three single-cell patch-clamp recordings and high-density microelectrode array (HD-MEA) measurements, in parallel. Our results demonstrate that the eANN outperforms all other algorithms across different dynamical regimes and provides a more accurate description of the underlying topological organization of the studied networks. We also provide a SHAP analysis of the trained eANN to understand which input features of the eANN contribute most to this superior performance. The eANN is a promising approach to improve connectivity inference from spike-train data.

## Introduction

Inferring the wiring diagram of complex neuronal circuits, their *connectomes*, has remained an important pillar in the quest to understand how individual neurons process information and how neuronal networks are organized [1]. Recent years have seen significant advances in connectome inference techniques enabling the study of fundamental principles of neuronal organization [2], and the intricate structure-function relationship of local synaptic connectivity [3, 4]. In particular, methods based on serial block-face electron microscopy (EM) [5], and virus-based circuit reconstruction [6], have paved the way to mapping out synaptic connections at unprecedented detail and scale. These methods significantly furthered our understanding of how circuit architecture relates to neuronal communication across scales, i.e., at the level of local connectivity, as well as, across different brain regions. The interest in linking connectomics and functional readouts has also been fuelled by the ever-increasing capabilities of large-scale electrophysiological recording technology for studying neuronal physiology *in vivo* [7] and *in vitro* [8, 9].

A large body of studies, including different species and brain regions, has started to provide insight into the specific connectivity patterns that individual neurons form to communicate. Common organizational motifs of synaptic connectivity include, for example, feedforward excitation, feedforward inhibition, as well as, feedback inhibition, and lateral inhibition [10]. In addition to these circuit motifs, a range of complex topological properties have been described [2], among them, a greater-than-random community structure [11, 12], the occurrence of specific triple-motifs among locally connected projection neurons [13, 14], a small-world [15] and rich-club organization [16], and highly-connected hub neurons [17]. Studies also found that the synaptic strength of local circuitry typically follows a heavy-tailed log-normal distribution, with few strong connections [18, 19]. Many of these synaptic wiring diagrams have been obtained through EM reconstruction in model organisms, such as *Caenorhabditis elegans* [20], *drosophila* [21], and *zebrafish* [22, 23], *and more recently, through reconstruction of small tissue samples of mouse* [4, 24], *macaque, and human cortex [25]*.

In addition to dense, EM-based reconstruction of neuronal circuits, which allows for the perhaps most comprehensive characterization of synaptic connectivity, important alternative circuit-mapping tools exist. The two most widely used techniques are viral retrograde and anterograde trans-synaptic labeling of neurons [6, 26], and whole-cell patch clamp recordings [27]. Patch-clamp recordings are the gold standard to infer synaptic function and have been widely applied to characterize synaptic connections among pre- and postsynaptic neurons, including the strength of their excitatory and inhibitory postsynaptic potentials (EPSPs/IPSPs), the time course of the postsynaptic responses, and the reliability of synaptic transmission. To assess if two neurons are monosynaptically connected, typically, whole-cell recordings in both cells are obtained, and spikes are induced via brief repetitive stimuli to measure the evoked EPSP/IPSP amplitudes and the direction of the connection(s) [13]. Whole-cell recordings from up to twelve simultaneously recorded neurons have also been used to study the organization of local connectivity in brain slices [3, 13]. Overall, the data obtained from patch-clamp recordings have provided essential information on the mechanisms underlying neuronal circuit computation, that, so far, could not be provided by EM-based reconstructions. This holds particularly for patch-clamp studies that were performed *in vivo* [28], which do not suffer from potential slicing artifacts [27]. Despite attempts to automate and scale-up patch clamping procedures [29], the throughput for connectivity studies has remained comparably low.

Besides inferring synaptic connectivity from intracellular recordings, there has been a surge in studies that used the statistical relationship of the activity among neurons as an indirect measure of neuronal coupling [30]. Spike train cross-correlograms (CCGs), for example, have been applied to estimate spike transmission or effective connectivity between defined neurons and/or specific brain regions [31–36]. To improve the performance of CCG-based circuit inference, several modifications have been suggested – such as, to take into account co-modulating background dynamics [37–39], to apply model-based timescale separation techniques [40, 41] or, more recently, to apply deep learning methods [42]. Still, inferring synaptic connectivity from the ongoing spiking activity of neurons remains highly challenging [43]. This holds true, in particular, if the strength of synaptic connectivity is weak [40], if the neuronal networks cannot be fully sampled [44], and if the used spike trains exhibit strong temporal periodicity, e.g., caused by correlated network bursts [39, 45]. Such burst activity may lead to high spike train synchronicity between two neurons that, however, are not synaptically connected [45]. This limits the interpretability of CCG-based methods but also holds for other algorithms used to infer interneuronal coupling in neuronal networks, recorded with either electrophysiological [30] or optical methods [46, 47]. It is important to be aware of these caveats when interpreting the topology of neuronal networks obtained from such activity-based connectivity-inference methods [48–52].

In this study, we introduce a workflow to benchmark algorithms that have been used to infer neuronal connectivity from large-scale extracellular recordings [36, 40, 53–56]. We, therefore, standardize the output of algorithms and compare statistically inferred connectivity estimates on synthetic ground-truth data sets and experimentally obtained connectivity labels. The first ground-truth data set was generated by stimulating *leaky integrate-and-fire* (LIF) neurons with empirical spike-train data and statistically defined noise [41]. These data allowed probing the effect of varying network dynamics and recording lengths on network reconstruction performance. In addition, we assessed the performance of connectivity-inference algorithms on in vitro data, where connectivity labels, where inferred from parallel whole-cell patch-clamp and *high-density microelectrode array* (HD-MEA) recordings obtained from primary neuronal cultures [57, 58]. Finally, we introduce an *ensemble artificial neuronal network* (eANN) and probe whether ensemble learning techniques [59] can improve today’s methodology to infer synaptic connectivity. We train the eANN on the standardized output of all implemented network-inference methods and demonstrate that knowledge about the shared output from these methods does indeed lead to gains in network reconstruction performance. To this end, we use a *SHapley Additive exPlanations* (SHAP) [60] analysis to better understand the relative contribution of individual input features to the superior eANN performance, and to visualize and interpret which methods were driving the eANN model to predict excitatory or inhibitory synaptic connections.

## Results

### A framework for the systematic comparison of activity-based connectivity inference algorithms

To compare inference methods and evaluate their performance, we standardized the connectivity inference task: Each inference method received the same spike train activity, i.e., the spike times and the corresponding unit IDs of a given neuronal network recording. These data consisted of either a network simulation (see Sec. S1; Fig. 1**A**), or an HD-MEA extracellular network recording obtained from primary cortical cultures (see Sec. S3A). Then, as output, each connectivity inference method provided two results: (i) a directed weighted graph, that indicated the connectivity strength between all nodes, and (ii), a matrix that contained the *connectivity scores* (CSs) for all connections. We will refer to the first output as the *weight graph* (W), and to the second output matrix as the *score graph* (S). Each edge of the inferred connectivity score graph S represents a CS.

**Fig 1.**
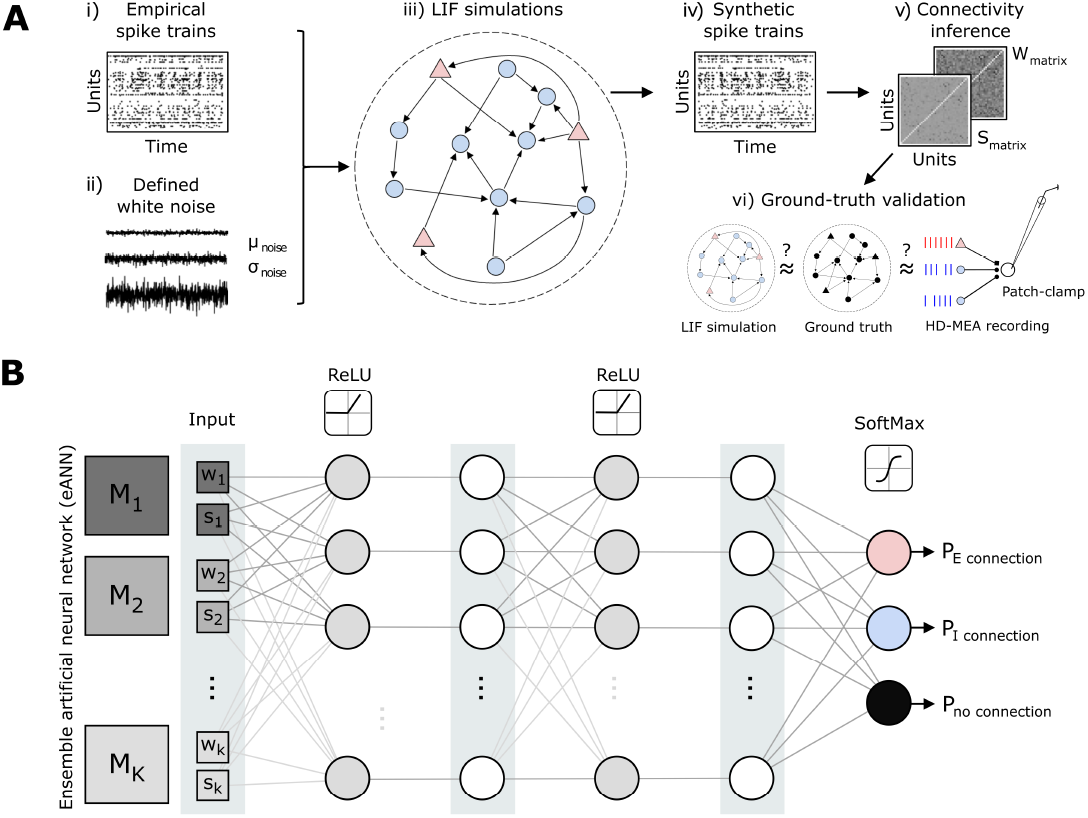
An ensemble artificial neural network to improve neuronal connectivity inference. **A** Schematic illustrating the developed analysis workflow to systematically compare statistically derived neuronal connectivity across inference algorithms and defined network dynamics. Empirical spike train data (i), obtained by high-density microelectrode array (HD-MEA) recordings from primary cortical cultures, and different types of white noise (ii) were used as input to a network (iii) of leaky integrate-and-fire (LIF) neurons (300 neurons, 50:50 excitatory (E) and inhibitory (I) neurons), adopted from previously reported work [41]. The *a priori* defined structure underlying the LIF network served as the first ground truth to compare established and new connectivity inference methods providing a score (*s*) and a weight (*w*) for each connection (v-vi). Moreover, connectivity-inference performance was also assessed on ground truth data obtained from parallel HD-MEA/patch-clamp recordings [58]. **B** Schematic depicting the architecture of the ensemble artificial neural network (eANN). The eANN receives as input the connectivity score *s* and weight *w* values from multiple established inference algorithms. Then the feed-forward network is trained. Finally, the eANN outputs probability values which indicate whether the connection is excitatory, inhibitory, or if there is no connection at all.

For a given putative connection between a presynaptic neuron *i* and postsynaptic neuron *j*, we defined a CS *s*_*i→j*_. The CS indicates the likelihood of a putative monosynaptic connection according to the respective inference method. Of note, the meaning of the CS depends on the method: For example, the CS could be a (log) probability, a negative (log) p-value, or the absolute value of a z-score (see Sec. S5 for the definition of CS and Table 1). As for the CS graph *S*, the interpretation of the weight graph *W* depends on the respective inference method, e.g., it might indicate an estimate of synaptic strength or how much information about the spike train *j* is contained in spike train *i*. For a given connection from neuron *i* to *j*, we define a weight *w*_*i→j*_.

**Table 1.** Overview on connectivity inference methods and abbreviations.

| Method name | Abbreviation | Description |
| --- | --- | --- |
| Coincidence index | CI | Measures the excess/deficit of coincidental firing compared to what would be expected by chance (shuffled data). Adpoted from [56] |
| Smoothed cross-correlogram | sCCG | Quantifies the temporal correlation between two spike trains; a smoothing kernel is used to discern background activity from synaptic interactions [37] |
| Directed spike time tiling coefficient | dSTTC | Quantifies the synchrony between two neurons by considering the temporal overlap and rate of spikes in a defined temporal window [53]. |
| Generalized linear model (GLM) cross-correlogram | GLMCC | Parameterized generalized linear model to facilitate synaptic connectivity inference from cross-correlograms. [40] |
| Transfer entropy | TE | Measures how knowledge about the spike history of one neuron reduces the uncertainty about the future spiking of another neuron [54,65] |
| Generalized linear model (GLM) point process | GLMPP | Generalized linear model to parametrize firing rate according to network history. Modified from [72] and combined with methods from [73]. |
| Ensemble artificial neural network | eANN | Neural network trained on CI, sCCG, dSTTC, GLMCC, TE and GLMPP to predict connectivity, as introduced in this study. |

We consider a putative connection between neuron *i* and *j* as present when the respective confidence score *s*_*i→j*_ is larger than a statistically defined threshold. This threshold, however, has to be defined by the experimenter. Connections below this defined threshold are regarded as unconnected pairs. As synaptic connectivity is generally assumed to be sparse, this task is highly unbalanced, i.e., we would expect that the unconnected pairs largely outnumber actual connections. Such imbalance poses a problem for accuracy measurements [61]. To compare network reconstruction performance across algorithms, and to take this imbalance into account, we here considered the averaged precision score (APS). We defined the APS as the integral of the precision-recall curve, which attains values between 1 (the best possible outcome) and 0 (the worst outcome). Since the APS is an aggregate statistic over all possible thresholds and some analytical questions do require thresholded graphs, we also considered the Matthews correlation coefficient (MCC) [62] as a second performance metric. See Supplementary Information S6 for details.

### Overview on connectivity inference algorithms

Historically, many studies have applied cross-correlograms (CCGs) to estimate putative mono-synaptic connections between neurons [31–33, 35–37]. In the present study, we considered three CCG-based connectivity inference methods: First, the *Coincidence Index* (CI), which integrates the CCG over a small synaptic window [56] and compares it to values obtained from jittered surrogate data (i.e., spike trains for which the short-latency synaptic relationships have been destroyed). The second method convolves CCGs with a partially hollow Gaussian kernel [37] and thereby allows separating slower background activity from faster synaptic interactions. We refer to this method as the *smoothed cross-correlograms* (sCCG) algorithm [36]. And finally, a third method, which fits a *generalized linear model* (GLM) to the CCGs [40]. As in the original publication [40], we refer to this method as the GLMCC algorithm. The GLM models the background spiking and the potential synaptic effect as two separate additive functions: The stronger the contribution of the synaptic effect is, the more likely a synaptic connection.

Several studies have applied information-theoretic measures to estimate neuronal connectivity and information flow between brain regions and individual cells [51, 52, 54, 63, 64]. Here, we utilized an efficient algorithmic implementation of *transfer entropy* (TE) [65] to infer connectivity from the discretized spike trains. TE has been used as a measure of information flow and quantifies – in this context – if information about the spike activity of neuron *j* improves the prediction of the activity of neuron *I* in addition to knowledge about the spiking history of neuron *i* alone [64].

We also implemented a modified, directed variant of the *spike time tiling coefficient* (dSTTC) [53]. Although originally not developed to quantify synaptic connectivity, but rather the synchronicity between pairs of spike-trains, it has recently gained a lot of popularity [66–68]. The dSTTC variant used here quantifies interneuronal coupling by estimating the proportion of spikes of two units that appear within a specific synaptic window. More specifically, dSTTC checks whether there is an access or scarcity of spikes of neuron *j* following the spikes of neuron *i* and spikes of neuron *i* that were elicited before those of neuron *j*.

Finally, the last inference method was based on a *generalized linear model for point-processes* (GLMPP) [55]. The GLMPP model allows the integration of multiple covariates, such as the past activity of the entire network, in an explanation of the observed spiking activity of a neuron *i*. Models of this class have been used in the past for inferring connectivity among partially observed neuronal populations [69].

Each inference method was adapted to the previously described analysis framework, i.e., each method provided a *score graph* S and a *weight graph* W. For more details on all implemented algorithms, and the applied modifications to fit them to our workflow, see *Supplementary Information* Sec. S5. An overview of all methods is provided in Tab. 1.

### Using an ensemble artificial neural network to improve connectivity inference from spike trains

Due to the great diversity of neuronal cell types and connectivity, and the inherent complexity of neuronal dynamics, it is likely that network-reconstruction algorithms perform better in some scenarios than in others [46, 70]. In this study, we introduce an algorithm that is based on an *ensemble artificial neural network* (eANN), and which can make predictions about synaptic connections after being trained on some training ground-truth simulation with the corresponding output of multiple inference methods (see previous section). Hence, one goal was to probe whether the collective input from traditional inference algorithms – as learned by the eANN – improves network reconstruction accuracy and robustness (Fig. 1).

To address this question, we first trained the eANN on connectivity-inference results obtained with the traditional algorithms from *leaky integrate-and-fire* (LIF) network simulations (see Sec. S1). For a putative connection between neurons *i* and *j*, the resulting eANN received the weight *w*_*i→j*_ and connectivity scores *s*_*i→j*_ of all six inference methods as input. The implemented eANN architecture consists of a feed-forward network [71] with two hidden layers with ten units each and a rectified linear unit (ReLU) non-linearity. The output layer has three units and a softmax non-linearity and indicated either an excitatory (E), an inhibitory (I), or no connection at all (Fig. 1**B**). Because we hypothesized that excitatory and inhibitory connections might be reflected differently by each method, we chose a multi-class classification setting. The model was then trained on different LIF simulations by minimizing the cross-entropy loss using the ground-truth labels in the training data. The final connectivity score *s* of the eANN was defined as the maximum softmax output for either an excitatory or inhibitory connection. If the CS for a connectivity pair *i→ j* exceeded a specific predefined threshold, the connection was deemed significant. The eANN method simply predicts if a connection exists (or not), without assigning a weight *w*. Predicting actual synaptic strength is expected to be a more challenging task, which we leave to future research. Note, that throughout this paper we train only the network once, and subsequent results are obtained by this network.

### Comparing network reconstruction performance across algorithms, network dynamics, cell types, and recording duration

To benchmark connectivity-inference methods, we first turned to LIF simulations (see Sec. S1), which were obtained similarly to an approach outlined in a previous study [41]. The spike train output of the LIF simulations resembled the subsequently used *in vitro* HD-MEA recordings. Each LIF network consisted of *N* = 300 neurons, equally split into excitatory and inhibitory neurons, and connected randomly with a probability of 0.05. The LIF network received two types of input: i) spike train data from an experimental recording to mimic realistic dynamics (connection probability to LIF neurons 0.1), and ii) white noise input. The white noise was varied, to map out a range of different dynamical regimes. As benchmarking datasets, we generated three different simulations: a high-bursting regime (burst rate: 1 Hz, average firing rate: 1.6 Hz, coefficient of variation: 0.3, see Fig. 2**A**), an intermediate-bursting regime (0.4 Hz, 1.1 Hz, 0.4), and a low-bursting regime (0.2 Hz, 1.3 Hz, 0.2). To test the robustness of our algorithms, we grouped each simulated network dataset into three subsets, each composed of 100 neurons (50 excitatory and 50 inhibitory neurons); each simulated dataset was 60 min long.

**Fig 2.**
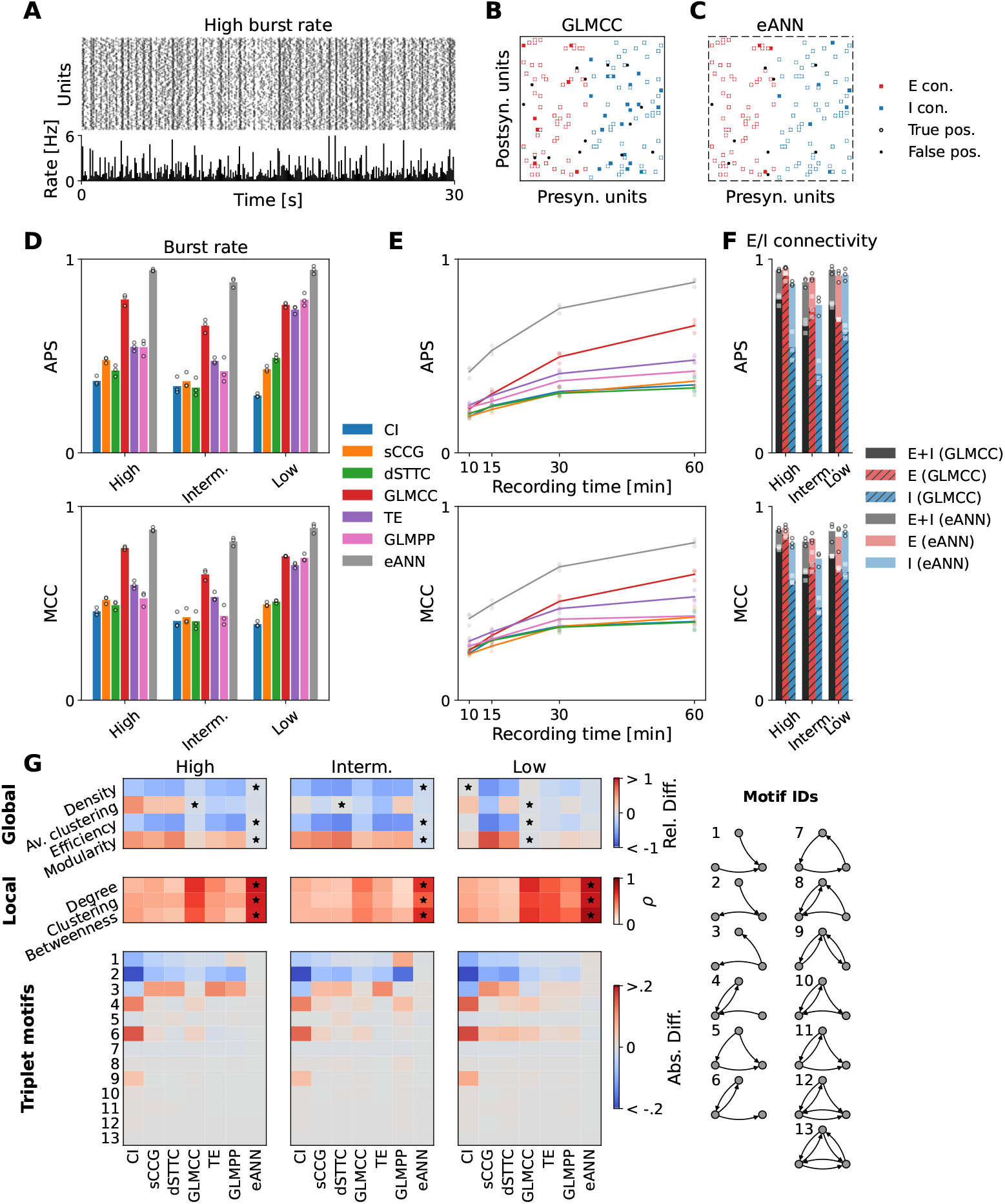
Reconstruction performance across algorithms, dynamics, cell types, and recording length. **A** Example raster plot (upper panel) and traces of binned population activity (lower panel; the number of spikes per second and neuron) of the *high-burst rate* condition. **B** Network reconstruction obtained from a subset of the data shown in **A**, exemplified for the GLMCC method [40]. Red and blue squares correspond to ground-truth excitatory and inhibitory synapses. White and black circles are predicted true positives and false positives. **C** Same as **B**, for the results obtained with the eANN approach, which generally improved the reconstruction performance. **D** The mean average precision score (APS, upper panel) and Matthews correlation coefficient (MCC, lower panel), estimated across all connections, obtained from all inference algorithms and the eANN across three different dynamical regimes. Dots depict the performance obtained on three different subnetworks of the same simulation. **E** Connectivity reconstruction performance (APS, MCC) as a function of recording time. Results indicated an improvement for longer recordings. Results in panel **E** are depicted for the intermediate dynamical regime. **F** APS and MCC for each type of connectivity, that is, excitatory (E, in red), inhibitory (I, in blue), combined (E+I, in black). Correspondingly, the performance gains achieved by the eANN are plotted in shades of red, blue, and black. **G** Quality of topological feature reconstruction for the inferred network across the three dynamical regimes. In the upper panel, the relative difference between four global features (network density, average clustering, and efficiency) is shown. The panel in the middle shows average Pearson correlation coefficients for the local/nodal metrics, comparing values obtained for the ground truth and inferred networks. Black stars indicate the method that performed best. The lower panels depict the absolute difference of triplet-motif frequencies between the ground truth and the inferred networks.

In Figure 2, we provide an overview of the benchmarking results for all implemented inference algorithms, including the eANN, across different activity regimes, excitatory and inhibitory connectivity types, as well as network inference calculations run on fewer data (subsets of 10, 15 or 30 min of the data; see subsampling analysis). Example activity for the simulated high-burst regime is illustrated in Fig. 2**A**. A network, reconstructed by two of the best-performing algorithms (GLMCC and eANN), is depicted in Fig. 2**B** and **C**; the average APS and MCC statistics across all methods are depicted in **D**-**E**); the reconstruction performance for excitatory and inhibitory connectivity is shown in panel Fig. 2**F**.

In line with previous research [43, 46], we found that the network reconstruction from the spiking activity was altered, if the provided spike train data contained periods of very synchronous activity. This became also apparent in the results depicted for the GLMPP and TE algorithm (Fig. 2**D** and **F**). While the performance of these algorithms was good for the low burst regime (e.g., TE: APS=0.74, MCC=0.70; recording duration: 60 min; connectivity thresholded at maximum MCC value), it deteriorated strongly for recordings for the high burst regime (TE: APS=0.55, MCC=0.59). Here, the inferred connectivity suffered from many false positives. The performance for the GLMCC method seemed less affected (GLMCC: APS=0.76 (low)/0.79 (high), MCC=0.74/0.79). The performance of the GLMCC algorithm for excitatory connections was fairly good overall (Fig. 2**B**; excitatory connections in red), but less robust for inhibitory connections (few false positives; Fig. 2**B** and **F**; inhibitory connections in blue). For all methods, reconstruction performance decreased for shorter recording lengths (Fig. 2**E**), again, in agreement with previous reports [40, 74]. Very similar performance profiles were also observed for most of the other inference algorithms (Fig. 2**D**-**E**).

Next, we probed the network-reconstruction performance of the eANN. Results indicated that the eANN outperformed all other inference methods – both, across all dynamical regimes (Fig. 2**D**), and when applied to shorter recordings, respectively fewer data (Fig. 2**E**). The most substantial improvements were observed for the intermediate-burst regime. Here, the average APS for the eANN was 0.88, i.e., a 35% improvement compared to the best-performing model (GLMCC: APS=0.65). Similarly, we found a 24% improvement in the MCC values for the eANN (eANN: MCC=0.81) compared to the GLMCC (eANN: MCC=0.65; see Fig. 2**D**). The main performance gains for this condition resulted from a reduction in false positive excitatory connections and an increase in true positive inhibitory connections (see Fig. 2**B**-**C**). For the temporal subsampling analysis (Fig. 2**E**), we compared the reconstruction performance of all algorithms on 10, 15, 30, and 60-minute subsets of the data. Although the eANN performance also decayed for shorter data segments, it was still significantly better compared to the other methods. Interestingly, the accuracy values of the best-performing traditional inference method, the GLMCC algorithm, degraded strongest on shorter recordings (lowest ASP/MCC values for spike train data below 30 minutes). See Fig. S1 for the precision-recall curves across all datasets and algorithms.

Finally, we probed how accurately the different algorithms could infer the global and local topological statistics of the simulated ground-truth networks. For the global metrics, we quantified the dissimilarity between the inferred networks *F*_inf_ and the ground-truth networks *F*_gt_ by their relative distance (*F*_inf_*− F*_gt_)*/F*_gt_. For the local topological metrics, we computed the Pearson correlation *ρ* between all nodal values of the inferred network and the nodal values of the ground-truth network; the reported values were averaged over three different networks. In Fig. 2 **G**, we report summary results for reconstructed graphs that were binarized with the best-performing MCC value. We found that many of the inferred connectivity metrics, including basic properties such as the network density, but also nodal features (e.g., the average clustering coefficient), differed substantially between algorithms and dynamical regimes. As expected from the results depicted in Fig. 2 **D**, most connectivity methods, and in particular the GLMCC algorithm, performed reasonably well in the low-burst regime. Here, we found only little relative differences between the global/local topology of the ground-truth networks and the topology of the inferred networks. For the intermediate and high-burst regimes, however, the estimates became worse. Here, the eANN excelled and outperformed all other methods. The average relative difference in global topological metrics for the eANN was 4%, 10%, and 6% for the low, intermediate, and high-burst regimes, respectively. For the second-best method, the GLMCC algorithm, the relative difference in global topological metrics were 2% (low), 27% (intermediate), and 13% (high-burst regime). For the local features, the eANN performed best on average (Pearson’s correlation: 0.74, 0.63, 0.81 for the high, intermediate, and low-burst regime); the performance of the GLMCC algorithm was still good, but lower on average: 0.62 (high), 0.44 (intermediate), and 0.65 (low-burst regime). The good reconstruction performance of the eANN algorithm became even more apparent for the inference of triplet motifs [18]. While many algorithms showed similar results in their estimates of the frequency of motifs, the eANN performed overall very well (Fig. 2 **D**). Of note, in Fig. S2, we also report the topological analyses for graphs binarized with a threshold obtained from jittered surrogate data (*α* = 0.01). As for the results presented here, we observed that the eANN showed a very robust performance.

While the eANN does not provide a prediction of the synaptic strength, we observe that its CS *s* is correlated with synaptic strength (see Fig. S8). Furthermore, we investigated the performance on a LIF network simulation with an 80/20 E/I connectivity in Fig S7, where we observed better overall performance of all connectivity inference methods with the eANN being the most performant method.

### SHAP analysis of eANN output

Next, we sought to understand in more detail which input features of the eANN contributed most to the improved reconstruction performance. We, therefore, investigated the eANN output with a SHAP analysis [60]. A SHAP analysis allows determining how much each of the connectivity methods (i.e., the input features) adds to the eANN predictions of excitatory or inhibitory synaptic connections. The SHAP values, which are calculated for each of the connectivity methods and each prediction, represent the contribution of the individual methods to the prediction task - and thereby explain party the inner workings of the eANN model. Figure 3 depicts the SHAP values for 1500 putative connections obtained from an eANN analysis performed on simulated spike trains in the intermediate-burst regime. In panel **A**, we depict SHAP values for example inhibitory and excitatory connections. The results indicate that approximately three features contributed most prominently to the decision of the eANN in favor of an excitatory (E) connection (upper panel), namely, the obtained weight of the GLMPP method and the connectivity score (CS) values of the TE and sCCG algorithms. For the inhibitory (I) connection (lower panel) mainly the weights of GLMPP and GLMCC methods contributed to the decision. In the figure, the contribution to the eANN output can be inferred from the length of the arrows, here depicted in green, respectively purple (Fig. 3**A**). While many features seemed to carry information about excitatory connections (Fig. 3 **B**, top panel) only about two features were predictive for inhibitory connections. However, the SHAP values for these features were quite variable. Interestingly, we observed that the features that yielded – on average – the highest SHAP values, were the same for excitatory and inhibitory connections (i.e., the weight estimates inferred by the GLMPP and GLMCC algorithms). However, we also found differences in how features contributed to detecting excitatory and inhibitory connections. For example, while the CS of the sCCG method contributed to the detection of excitatory connections, it contained only little information for the inference of inhibitory connections. To illustrate how single features contribute to the classification of excitatory/inhibitory connections, we depict the exact SHAP values of the two top-ranked metrics (GLMPP and GLMCC) as a function of their feature values (Fig. 3**C**): Results indicate that the two features were mainly negative for inhibitory connections (depicted in blue) and positive for excitatory connections (depicted in red); high SHAP values often correlated with large absolute feature values. This held, in particular, for the negative weight estimates of the GLMPP, indicating that this feature may contribute strongly to the identification of inhibitory connections. It is noteworthy, that none of these features alone was sufficient to separate actual connections from unconnected pairs. We propose this underscores that the eANN approach is advantageous for the reliable inference of connectivity from spike train data. Additionally, we investigate the correlations among input features in Fig. S11, where we observe that for excitatory connections features are much stronger correlated than for inhibitory connections. This yields evidence, that different input methods carry different kinds of information, and that the eANN can leverage this fact best for inhibitory connections yielding a strong performance boost for these.

**Fig 3.**
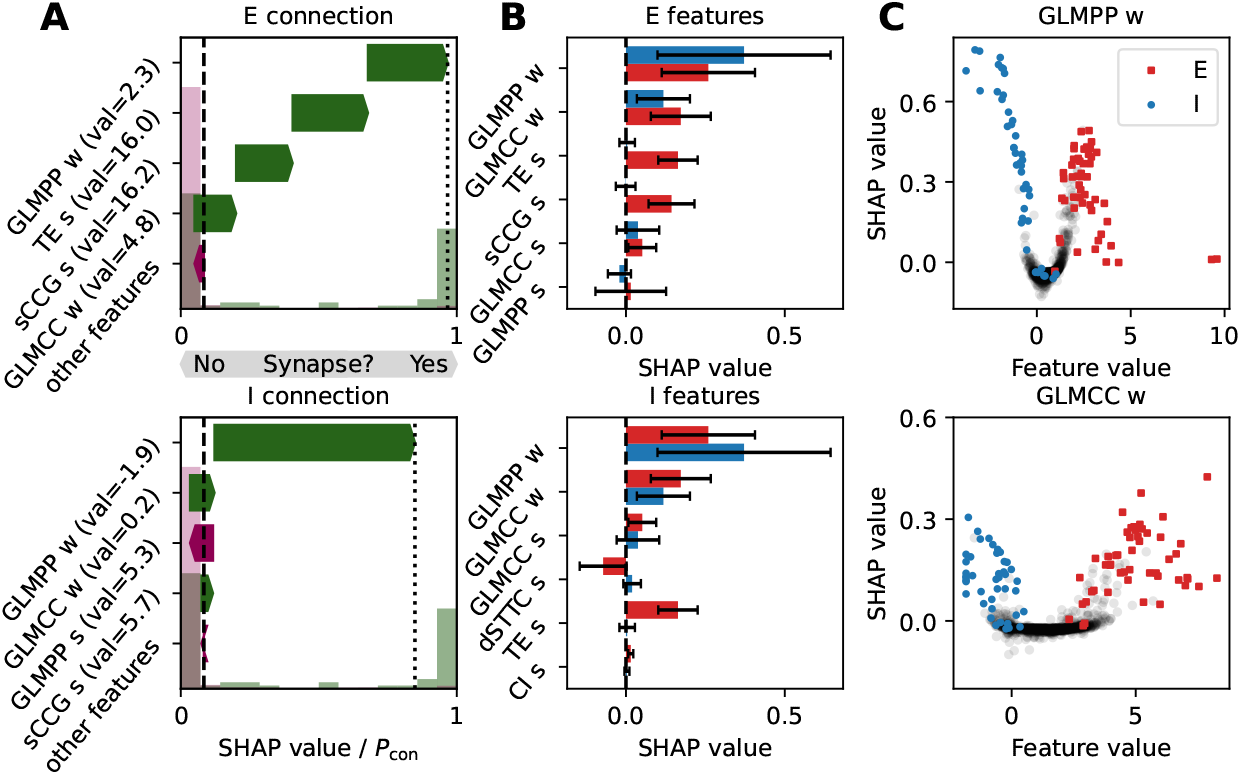
SHAP analysis on eANN output. **A** SHAP values for example an excitatory (E, top panel) and an inhibitory connection (I, bottom panel), ranked according to their importance. The length of the arrows denotes the approximated feature contribution to the eANN output, as estimated by the SHAP method [60]. The y-axis shows the corresponding features with the feature value in brackets. Green arrows pointing to the right indicate that this feature was informative in the process of determining an E/I connection; purple arrows pointing to the left indicate the opposite. Normalized histograms, plotted in light green and purple in the background, show the eANN output for unconnected pairs and connections respectively. The dashed line is the average eANN output for all connections in the dataset. **B** The features with the highest average SHAP values for excitatory (top) and inhibitory connections (bottom). Red and blue bars denote the average SHAP values for excitatory and inhibitory connections, respectively. The error bars denote the standard deviation, calculated over all excitatory/inhibitory connections present in the dataset. **C** SHAP values as a function of feature value for the two top features depicted in panel **B**, namely GLMPP w, and GLMCC w. Black dots indicate unconnected pairs.

### Application to HD-MEA/patch-clamp data

Next, we applied the developed inference pipeline to experimental data, obtained through parallel HD-MEA network and whole-cell patch-clamp recording in *in vitro* neuronal cultures (dataset 1; for details on the data see [58]; Fig. 4; n=3 patched cells; culture age: DIV16-18; see Sec S3E for details). Briefly, following a long HD-MEA baseline *∼* 3 h long recording for spike-based connectivity inference, we transferred neuronal cultures to a setup that enabled simultaneous HD-MEA/patch-clamp recordings. Here, single neurons were patched on the HD-MEA and, in parallel, recorded extracellularly on the HD-MEA. Two distinct paired recordings were obtained from each patched cell. First, we recorded spontaneous spiking of the patched cell in whole-cell current-clamp mode in addition to simultaneously recording extracellular HD-MEA signals. The obtained data then allowed to perform spike-triggered averaging (Fig. 4 **A**-**E**) and to infer the *electrical footprint* (EF) of the patched neuron. The EF is the extracellular profile of a neuron on the HD-MEA (Fig. 4 **D** and **E**).

**Fig 4.**
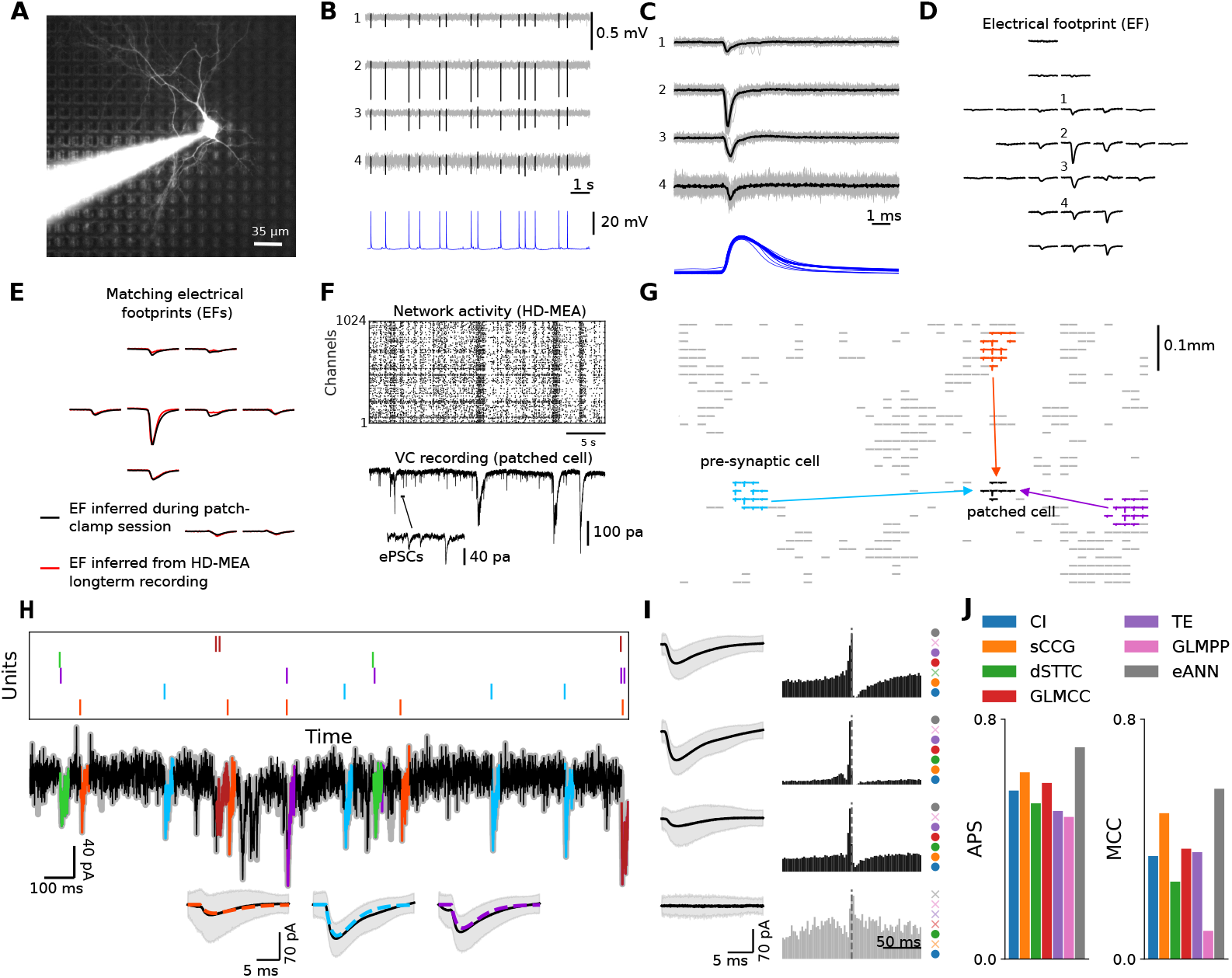
Validating synaptic connectivity inference with parallel HD-MEA and patch-clamp recordings. **A** Panel depicts a single patched neuron on the HD-MEA, including the patch pipette. The neuron was labeled with Alexa Fluor 594 through the patch pipette; HD-MEA electrodes can be seen in the background. **B** Example recording of a patched cell and some of its extracellular signals, obtained with HD-MEA network recordings (spikes are depicted in black) that were conducted in parallel to the patch-clamping (lower panel, in blue; whole-cell patch-clamp mode). HD-MEA and patch-clamp signals were aligned **C**, and allowed for inferring the exact location and electrical footprint (EF) of patched neurons on the HD-MEA **D**. To infer the putative pre-synaptic connectivity of individual (patched) neurons, we first performed long HD-MEA network recordings. Next, we applied a post-processing step to match the EF of the patched cell, with the EF templates obtained via spike sorting of the HD-MEA network recordings. The panel **E** depicts the overlap of the EF obtained during the patch session (in black) and the EF obtained from spike-sorted data (in red). **F** Extracellular network recording (upper panel, raster plot) and simultaneous intracellular current signal (lower panel), obtained from a whole-cell voltage-clamp (VC) recording. VC recordings were used to measure the excitatory/inhibitory postsynaptic currents (ePSCs/iPSCs) in the patched cell, and their occurrence to the activity obtained from spike-sorted HD-MEA data. Panel **G** depicts three example connections of the patched cell (EF in black) to three presynaptic neurons; the EFs of these cells on the HD-MEA are colored in light blue, orange, and purple. **H** IC model fit to patch-clamp recording. Spikes of identified presynaptic neurons are shown on top, aligned with the recorded currents of the patched neuron. PSCs of three units are shown as insets, with the average depicted in black and the fitted model PSC displayed in different colors. The shaded area is the 5 *−* 95% quantile. PSCs of four cells, three connected pre-synaptic neurons, and one unconnected neuron, are shown along with their corresponding CCGs. Note, that a high-chloride internal patch-clamp solution was used to simultaneously measure the synaptic activity of excitatory and inhibitory presynaptic cells [58]. As a result, as depicted, inhibitory input currents also have a negative polarity. On the right side of the CCG, a colored circle indicates whether the respective connectivity method found a connection or not (cross). **J** Panel depicts the performance of the different connectivity methods averaged over the three patched neurons.

To study the synaptic input to the individual postsynaptic cells during ongoing spontaneous activity, we recorded neurons in voltage-clamp mode (Fig. 4 **F**). As previously reported [57, 58], combining the spontaneous/evoked electrical activity of neuronal networks, recorded on the HD-MEA (Fig. 4 **F**), with the measured postsynaptic currents (PSCs) in patched neurons, allowed for reliably reconstructing connectivity to putative presynaptic partner cells. To estimate the effect of extracellularly recorded neurons on the patched cell, and most importantly, to infer synaptic connections between the patched neuron and its putative presynaptic network, we developed a regression-based connectivity inference method (similar to [75], see Sec. S2 for details). The method was first benchmarked on *in silico* data; an overview of the obtained performance results is provided in Fig. S5. Our *in silico* results indicated that the developed method can robustly detect synaptic connections (high recall), with only very few false positives (high precision). Although the performance deteriorated for low-rate conditions and more correlated spiking, the method proved very reliable for a wide range of parameters.

Next, we applied the developed intracellular connectivity inference method to three patch-clamp recordings obtained from a previously published dataset [58], and identified 26 putative synaptic connections among 131 possible combinations. In Fig. 4**H** we show an example fit with three PSC cutouts of three identified connections in an example recording; Fig. 4**I** depicts the CCGs and PSCs of three connections found in the recording and an unconnected pair. For an overview of all PSCs and corresponding pair-wise CCGs of identified connections, as well as for information on which algorithms succeeded in detecting the connection, please see Fig. S6.

With this labeled experimental data at hand, we then evaluated the performance of the existing activity-based inference methods, and the eANN, on 3-h-long HD-MEA baseline recordings. The eANN is the same as for Fig. 2 and was not re-trained for these analyses. For the *in vitro* data, we see similar performance for the six input methods in terms of APS, but varying performance for MCC (see Fig. 4**J**). The GLMCC algorithm (MCC=0.37), the sCCG (MCC=0.49), and the TE algorithm (MCC=0.36) performed best among the input methods. The superior performance of GLMCC originates rather from higher precision than recall (recall=14 correctly classified connections of 20 connections; precision=14 correctly classified connections among 39 identified synapses). In comparison, the sCCG method (recall=23*/*26; precision=23*/*53) and the TE algorithm (recall=16*/*26; precision=16*/*38) both yielded similar or higher recall, at the expense of lower precision. As for the analysis of simulated ground truth, we found that the eANN (MCC=0.56) outperformed the other methods in terms of both MCC and APS. Also here the dominating factor was, that the eANN yielded much higher precision at inferior recall (recall=17*/*26; precision=17*/*26). Considering these numbers, in particular, for recall and precision, we found that this performance increase was mainly due to more accurate predictions (higher precision), at the cost of not identifying some true connections (lower recall). In summary, our results indicate that connectivity inferred by the eANN method is more precise than the results obtained with any of the other inference methods.

### Characterizing in vitro neuronal network connectivity and topology

Finally, we applied the eANN pipeline to a second dataset of HD-MEA network recordings, obtained from primary cortical cultures (dataset 2; n=6 cultures; recording duration: 1 h; culture age: DIV14). The main goal of this analysis was to compare how connectivity and topological properties varied across the implemented inference methods (Fig. 5). As before, all network recordings were first spike-sorted and underwent several quality-control steps to ensure that connections were estimated on sufficient activity (Fig. 5**A**, see Sec. S3D). To reduce between-network variability, we only sampled 100 units from each network. Comparing the overall distribution of empirical eANN scores (Fig. 5**B**, depicted in black) to the corresponding surrogate distribution of eANN output values (depicted in yellow) indicated that the spike-train jittering effectively destroyed short-latency synchronization between individual neurons. The distribution of experimentally inferred values showed a clear peak in the eANN weight distribution that distinguished putative synaptic connections from unconnected pairs (Fig. 5**B**). Next, we calculated the consensus distribution (Fig. 5**C**), i.e., the overlap between all significant connections, obtained by the six traditional inference methods and the network inferred by the eANN (threshold value: *α*=0.01). The results indicated that most eANN edges were found by five of the six inference methods (32.6%). Interestingly, some edges were found by the eANN method, but not by the other methods at the selected threshold (edges in zero bin: 18%). This finding indicates that some eANN edges could not exclusively be explained by the overlap across all inference methods, but that there is added value by the eANN algorithm at this threshold. The network density decreased with smaller *α*-values (Fig. 5**B**; *α*-values: 0.05, 0.01, 0.005 and 0.001), and varied significantly as a function of inference algorithms (the repeated measures’ analysis of variance (ANOVA) for a *α* threshold value of 0.01 was: F(6,30)=189.77, p=6.5659e-10; p-value with Greenhouse-Geisser adjustment); the network density of all graphs was sparse (e.g., for a threshold of *α*=0.01, the network density varied between 1-9%; Fig. 5**B** and **G**), in line with previous reports [18, 76]. All implemented algorithms reconstructed networks that showed a clear decay in connection probability as a function of interneuronal distance (Fig. 5**F**).

**Fig 5.**
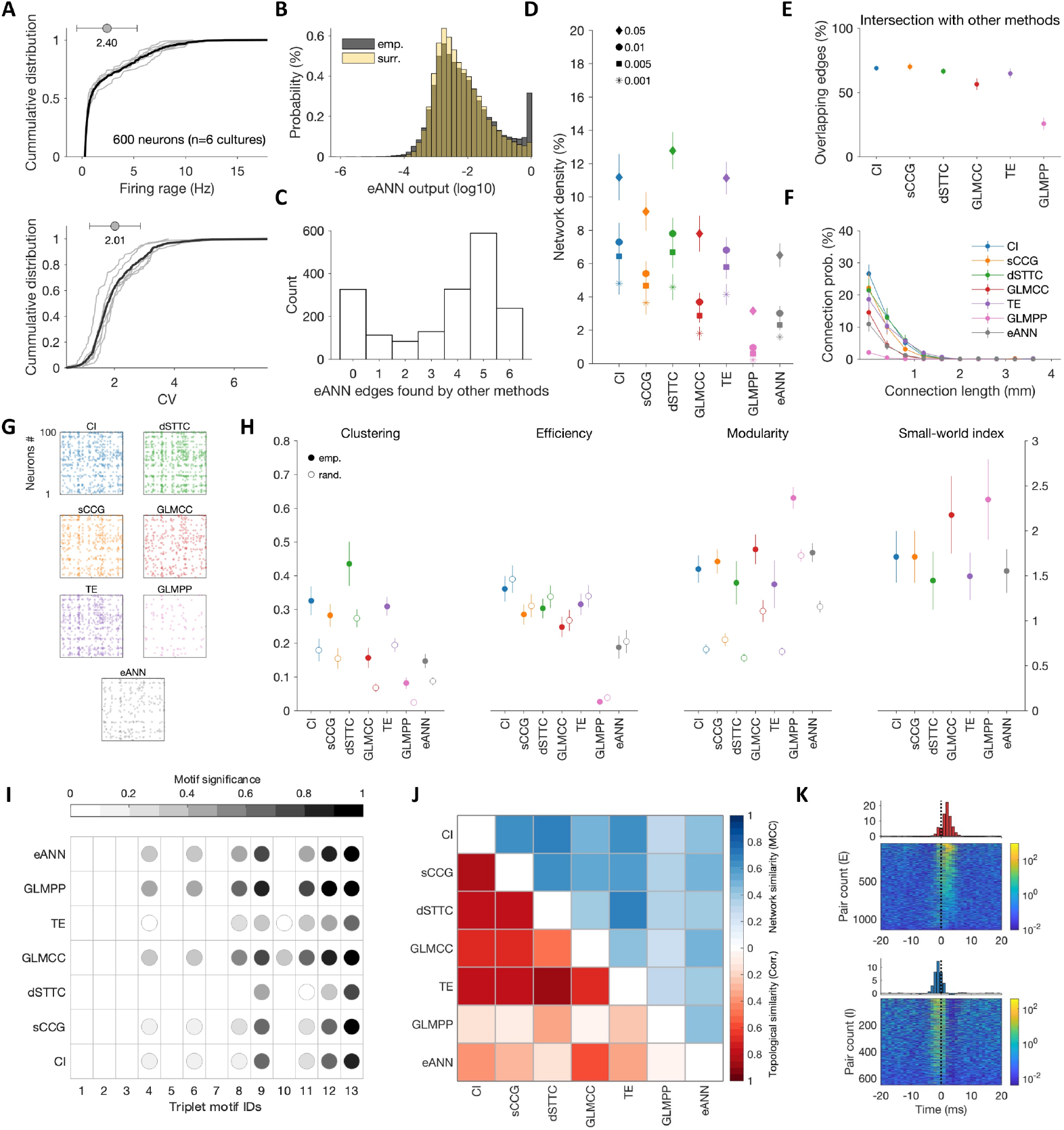
Characterizing the connectivity of in vitro neuronal networks. **A** Firing rate cumulative density function (CDF) of 600 spike-sorted units from HD-MEA recordings obtained from primary cortical cultures (top panel; n=6 cultures; 100 randomly selected units per network; recording duration: 1 h; culture age: DIV14); and CDF of inter-spike interval coefficients of variation (CV) for the same cultures (lower panel). **B** Network density as a function of threshold values (corresponding to *α*: 0.05, 0.01, 0.05, and 0.001) across all inference methods. *α* threshold values were derived from surrogate connectivity estimates (temporally jittered spike trains). Network density decreased with smaller *α*-values, and varied significantly across methods. **C** Overall distribution of eANN weights of empirical networks (in gray; values are depicted in logarithmic space) and overlaid with the corresponding distribution of eANN values inferred from surrogate networks (in yellow). The distribution of experimentally inferred values demonstrates a clear peak in the eANN weight distribution that distinguishes putative synaptic connections from unconnected pairs. **D** Intersection of significant eANN connections with those of all other connectivity inference methods. Panels D-K depict network graphs thresholded with *α* = 0.01. **E** Significant eANN edges cannot exclusively be explained by the overlap across all inference methods. Panel E depicts the consensus distribution, i.e., the overlap across the six inference methods for all significant eANN edges; most eANN edges were found by five of the other methods. About 20% of edges were found by the eANN, but not by the other methods at the selected threshold (see zero bin). **F** Inferred connectivity decayed with interneuronal distance (*α* = 0.01), and the likelihood of long-range connections (*>* 1 mm) was very low. **G** Example connectivity matrices for one culture inferred with all seven inference methods. **H** Topology of inferred in vitro networks differed significantly across inference methods (*α* = 0.01; filled circles: empirical data; empty circles: randomized surrogate networks). **I** All inference methods yielded an over-representation of triplet motifs (see Fig. 2 for motif ID legend) but with slight differences across methods. Dots depict motif IDs that occurred significantly more frequently than re-wired surrogate networks (FDR corrected *α* of 0.001); the color indicates the relative mean difference in their occurrence. **J** Lower triangle (red color scale): Topological similarity, calculated by pairwise Pearson correlation coefficients across the topological metrics shown in **H** and **I** across all inference methods. Upper triangle (blue color scale): Network similarity, quantified by pairwise MCCs across all adjacency matrices. **K** Display of putative excitatory (E) and inhibitory (I) connectivity for all edges inferred by the eANN. The top panel depicts the baseline corrected CCGs for the eANN edges with positive spike transmission probability (STP), i.e., excitatory connections. The average CCG is depicted in the red bar graph at the top. Correspondingly, the bottom panel depicts the putative inhibitory connectivity among the eANN inferred edges with negative STP. Note, the logarithmic color scale of pair counts. The excitatory and inhibitory connections are sorted according to their positive/negative STP values.

Furthermore, we found that topological properties of network segregation and integration differed as a function of the inference method (clustering: F(6,30)=102.16, p=1.67e-7; efficiency: F(6,30)=222.3, p=5.29e-14; modularity: F(6,30)=40.245, p=3.23e-5; small-world-index: F(6,30)=7.28, p=0.004; all p-values with Greenhouse-Geisser adjustment). Despite the reported differences, all algorithms implied that networks possessed a non-random modular, small-world organization (small-world-index *>* 1); a topological analysis with proportionally thresholded networks is provided in Supplemental Material (Fig. S3).

The inferred *in vitro* cortical networks also featured a significant over-representation of some triplet motifs [77] (Fig. 5**I**). The number of motifs that were significantly over-represented in the empirical networks varied between methods (e.g., four motifs for dSTTC and eight motifs for the GLMCC graphs), but all implemented algorithms suggested that motifs 9, 11, 12, and 13 were over-represented compared to an appropriate surrogate network (threshold: *α*=0.01; motif significance *α*-value=0.001; p-values were corrected for multiple comparisons across motif IDs and algorithms; Fig. 5**I**). Despite strong differences in the frequency of specific motifs between inference methods, there was a high resemblance across networks within each method (see Fig. S4). Results obtained from the topology and motif analyses indicated a higher similarity (correlation coefficient) between the output of the eANN and the GLMCC algorithm compared to the other methods (Fig. 5**J**, lower triangle). On average, topological properties and motif frequency values were more similar for the CI, sCCG, dSTTC, GLMCC, and TE algorithms – as compared to GLMPP and eANN. This trend is reflected in the pairwise Matthews correlation coefficient (MCC; Fig. 5**J**, upper triangle).

Finally, and to further supplement the topological properties, we also assessed the functional connectivity properties of eANN-inferred networks. We, therefore, calculated the *spike transmission probability* (STP) of significant eANN connections (threshold: *α*=0.01) using an STP-variant, that has been adopted and improved to better infer STPs on recurrent/bursty spike activity [39]. A positive STP value was taken as an indicator for an excitatory connection and a negative STP as an indicator for an inhibitory connection. Across all 1794 connections (2.99% overall network density), as inferred by the eANN algorithm (network threshold *α*-value: 0.01), 63.7% of connections were excitatory and 36.3% were inhibitory (Fig. 5**K**). Inspection of the CCGs indicated that many of the observed inhibitory connections might participate in inhibitory feedback motifs. Overall, these results underscore that the eANN algorithm was able to pick up connections of both excitatory and inhibitory types, supporting our previous modeling results.

## Discussion

The present study demonstrates that inference of synaptic connectivity from extracellular spike train dynamics can be improved by the application of an *ensemble artificial neural network* (eANN). By benchmarking the eANN to more traditional connectivity-inference algorithms in a standardized analytical framework, we report a superior reconstruction performance for the eANN, that persisted across different dynamical regimes and recording durations. We find that the eANN did also provide a more accurate reconstruction of the type of connectivity (excitatory or inhibitory), and better estimates for the global and local topology of networks *in silico*. Results derived from a SHAP analysis allowed for further validation of the specific contributions of algorithms to the eANN performance. Importantly, we found improvements in network reconstruction for both simulated and experimental ground-truth datasets. Such generalizability indicates that the developed method leads to more accurate and robust connectivity inference in datasets for which knowledge of the underlying synaptic connections is not available.

### The challenge to infer synaptic connectivity from spike trains

The extent to which synaptic connectivity and causal relationships between neurons can be studied, based on spike train dynamics alone, remains a matter of active debate. Although there have been attempts with small well-established circuits [78, 79], this endeavor has proven challenging for many reasons [43, 80]. In line with previous work [43, 46], the results reported in this study underscore, that there are indeed limits to activity-based network reconstruction (Fig. 2**D**). Using simulations to parametrically model a range of dynamical regimes – we found, as expected, significant performance alterations for most traditional algorithms once spike trains showed stronger temporal periodicity. With few notable exceptions, such as the GLMCC algorithm, reconstruction performance, quantified by the averaged precision score (APS) and Matthews correlation coefficient (MCC), worsened significantly for data with more prominent network burst-activity (Fig. 2**D**).

Among the many possible reasons for such performance, alterations are false positive connections generated through, for example, common inputs, poly-synaptic connections, or short-term synaptic dynamics [43, 45]. The accuracy to infer a synaptic connection from activity also crucially depends on the number of available spikes and synaptic strengths [74]. Weak connections and/or availability of only a few spikes drastically increase the likelihood of missing connections [81]. Moreover, insufficient coverage of the network may further alter the reconstruction performance of algorithms [82]. A combination of these factors most likely explains the observed imperfect inference performance, even for simulated recordings with less correlated population activity (Fig. 2**D-F**). Moreover, it should be noted, that the *a priori* conceptual assumption of what will be considered a connection, the choice of parameters for each method, and the criteria that are applied to falsify the existence of such connections (e.g., via surrogate generation) are likely equally important. Although previous studies have attempted to compare different connectivity reconstruction algorithms [39, 42, 70], most of them were less unified conceptually and are difficult to compare to the present study. Moreover, these studies used a variety of ground-truth data sets, including simulated calcium imaging data, which suffers from low temporal resolution [46, 47].

### eANN improves neuronal network reconstruction

Although ensemble methods have been proposed in the past [59], and used for the study of large-scale brain networks [83, 84], to the best of our knowledge, the present study is the first to demonstrate that such methods can be generalized to cellular spike-train data. More importantly, our results indicate that applying the eANN to such data allowed for significant performance improvements in the reconstruction of synaptic connectivity (Fig. 2; Fig. S1 and S2). Using a simple feed-forward network architecture [71], and training the eANN on the standardized output of several existing connectivity-inference methods [36, 40, 53–56], we demonstrate that excitatory and inhibitory connectivity, and the topology of neuronal networks, can be reliably inferred from simulated as well as empirical spike-train data (Fig. 2 and 4). Overall, the eANN outperformed all traditional methods (Fig. 2**D-G**), and the performance gains were relatively invariant to different dynamical regimes (Fig. 2**D**) and even persisted on fewer data (Fig. 2**E**). Our results also revealed that the eANN integrates aspects of different inference methods for its connectivity prediction (Fig. 5**C**), and that high eANN output values were absent, if the millisecond spike-timing information was disrupted, e.g., by jittering the spike trains (Fig. 5**B**). While understanding the performance of neural networks is typically challenging, insights gained from a SHAP analysis indicated that the trained eANN assigned varying levels of importance to different input features, i.e., some inputs were more informative than others (Fig. 3). Interestingly, this held particularly true for the detection of inhibitory connections (Fig. 2**F**). Here, the GLMPP algorithm seemed to convey valuable input (Fig. 3**B**, lower panel), a method that performed less well in the detection of excitatory connections. Overall, results for different connectivity types, network topology, and different network dynamics, as calculated with eANN-inferred graphs, were robust across different threshold definitions (Fig. 2, Fig. S1, and Fig. S2). We hypothesize, that the observed robustness could be attributed to the initial inference methods being designed to be somewhat invariant to differences in firing rates or baseline neuronal activity. Applied to HD-MEA network recordings, eANN-derived connectivity was most similar to graphs inferred by the GLMCC algorithm (Fig. 5**J** and Fig. S10). Despite these positive results, it should be noted, that the eANN did not result in perfect reconstruction performance. Thus, future studies should probe, whether considering additional features, more completely sampled recordings, or longer recordings could improve network reconstruction even further. Moreover, it would be interesting to probe, if inference performance improves, if data-driven features obtained from other neural network methods, such as the CoNNECT method [42], are integrated into the eANN.

### In vitro neuronal networks show complex topologies

Our results on the putative synaptic connectivity of *in vitro* developing primary neuronal networks, obtained by HD-MEA network recordings, and inferred by the eANN and traditional inference methods, are largely in line with previous reports [48, 49, 51, 52, 63, 74]. All algorithms indicated that connectivity was locally clustered, overall sparse (connection probability below 15%; Fig. 5**D**), and that the probability of connections decreased as a function of interneuronal distance (Fig. 5**F**). These results are also in agreement with previous patch-clamp work [13, 76]. Yet, it should be noted, that the effect of the inference method on connectivity and topology was considerable (Fig. 5**D**). For some topological measures, such as the clustering coefficient (Fig. 5**H**), the between-method differences exceeded the differences observed between the empirical and the randomized surrogate networks. For example, the clustering results, calculated on graphs inferred by the eANN and GLMPP methods, were significantly lower than the values of the randomized surrogate networks of some other methods (e.g., CI, TE, and dSTTC). Still, all networks comprised a modular small-world structure (Fig. 5**H**). Some subtle differences in the triplet-motif statistics (Fig. 5**I**) were observed in comparison to previous reports [63], but overall, these were not pronounced, and many of the over-represented motif structures have been reported by whole-cell recordings in slices [13, 18].

### Limitations

Several important limitations should be considered when interpreting the results presented in our study:

First, the analyzed spike-train data were obtained from dissociated primary rodent cortices cultured *in vitro*. Although such model systems have been used extensively to study neuronal physiology at the cellular level [8, 10], and have, thereby, provided fundamental insights into the mechanisms of underlying synapse formation and function, there are limits as to how insights obtained from *in vitro* connectivity can be translated to *in vivo* [85]. Future studies should, therefore, train and apply the developed eANN pipeline also to *in vivo* spike-train data – ideally, recorded in well-defined brain subsystems, where inferred connectivity can be linked to structural connections established with other methods [86].

The results presented in Fig. 5 were estimated on DIV14 neuronal networks, a time point where GABAergic signaling may be still immature [87, 88]. Although early during development, our results indicated the presence of some inhibitory connections among the recorded neurons (Fig. 5**K**). The relatively young age of these cultures, however, should be considered when interpreting the inferred network density. Likely, a significant proportion of synapses is still ‘silent’ at that time, and that some relevant receptors are not yet fully expressed [89]. Also, the cell-plating density of neuronal cultures, respectively the size of networks, affect overall synaptic strengths [76], and hence will impact the ability to infer synaptic connections from activity [90].

A common limitation, shared by all methods in this study, is, that they approximate connectivity from a network that is incompletely sampled. Such subsampling has been shown to lead to altered network reconstruction performance [43, 82, 90]. This holds also true for more complex inference algorithms that can take into account the past activity of the sampled network [54, 69]. Hence, the inferred connectivity may exhibit spurious false positives due to common unobserved input that cannot be explained away [43, 91]. This is also true for the experimental ground-truth data in our study. Live-cell imaging with calcium or voltage sensors could be applied to improve coverage [46, 70, 92], and future studies should compare how network statistics change as a function of network coverage.

We note, that also the empirical ground-truth data used in this study could be further improved [58]. Applying targeted, electrical stimulation to defined pre-synaptic neurons would allow for stronger claims about the found connections and their interneuronal causal effects [57]. Moreover, it would help to have access to the neuritic morphology of some parallel recorded neurons on the HD-MEA to better link the inferred connections to axonal/dendritic morphology and neuritic overlap. Such additional structural insights could also help to remove false positive connections.

## Conclusion

In sum, this study presents an analysis workflow to systematically assess algorithms that have been applied to the inference of neuronal connectivity from large-scale extracellular recordings. To this end, we utilized simulated and experimental ground-truth data, and compared statistically inferred connectivity across a range of different conditions in a standardized manner. Moreover, we introduced an *ensemble artificial neural network* (eANN), that can integrate the output of multiple inference algorithms, and probed whether this would lead to improved network-reconstruction performance. Our results demonstrate that inference performance can be significantly improved by the eANN, and that the obtained network reconstruction represents more than just the sum of all methods.

## Methods and Materials

Here, we describe the architecture and training scheme of the introduced *ensemble artificial neural network* eANN. All methodological details on the HD-MEA and patch-clamp experiments, the LIF simulations, the developed model to infer synaptic connections from parallel recordings, details on spike-time based connectivity inference methods, the *in vitro* culturing, the data preprocessing, as well as the performed topological analyses can be found in the *Supplemental Information*.

### Architecture of the ensemble artificial neural network

The introduced *ensemble artificial neural network* (eANN) is a feed-forward network, that takes the weight *W* and the CS *S* matrices of the six implemented connectivity methods (CI, sCCG, GLMCC, TE, dSTTC, and GLMPP) as input and provides as output the probabilities of whether the input belongs to an excitatory or an inhibitory connection, or whether the neurons are not connected at all (no connection). The eANN consists of two hidden layers with 10 units each and rectified linear units (ReLU) as non-linearity. The last layer is passed through a soft-max function, to normalize the output. We trained the model on several LIF neuronal network simulations subject to different inputs (see below). The trained network is the final eANN, which provides predictions based on the aggregated outputs of the other connectivity inference methods. The CS of the eANN was calculated as *s*_*i→j*_ = max(*p*_E_, *p*_I_), where *p*_E_ and *p*_I_ were the eANN’s predicted likelihoods for an excitatory or inhibitory connection, respectively.

### Training the ensemble artificial neural network

We simulated 25 different LIF networks to generate training data for the eANN. To obtain the activity of distinct neuronal network dynamics, we used five different noise configurations *µ*_noise_, *σ*_noise_ (see Table S1) and generated five networks with random connectivity for each configuration (connection probability: 0.05). Next, we simulated 1h of LIF spiking activity as previously described (see Sec. S1A). The experimental spiking input to the LIF network was always the same for the different training simulations. However, different input activity was used for testing the eANN (i.e., the data reported in Fig. 2). To train the network, we first obtained the connectivity output of all methods (CI, sCCG, GLMCC, TE, dSTTC, and GLMPP). Then, we took all synaptic connections, which make up 10% of the training set. For the remaining 90% of connections, we selected randomly unconnected pairs as negative examples. The CS and weight provided by the traditional methods constituted the input variables. The training labels were set to 0 (no connection), 1 (excitatory connection), and 2 (inhibitory connection), and the eANN was trained by minimizing the cross-entropy loss on these data. For the predicting connections on a new dataset, we again first applied the original inference methods and then provided their aggregated result as input to the eANN. We noted that the compact network architecture prevented from overfitting and ensured rapid convergence during training S9.

## Data, Materials, and Software Availability

The `PYTHON` code for all connectivity inference methods and the spike train datasets will be available at https://github.com/christiando/spycon. Raw data are available from the corresponding authors upon reasonable request due to their large size. Connectivity inference algorithms all ran on a cluster node, with 30 CPU cores (AMD EPYC) and 8-12GB RAM.

## S1 In silico simulations of neuronal networks

### S1A Leaky integrate-and-fire simulations

To test and compare network reconstruction performance across different algorithms, we adopted and modified the leaky integrate-and-fire (IF) network model approach proposed by Ren et al. [41]. The LIF simulations of the present study were performed with Brian2 [93]. The simulated network consisted of 300 neurons, balanced with 150 excitatory neurons and 150 inhibitory neurons. The differential equations of the IF model are

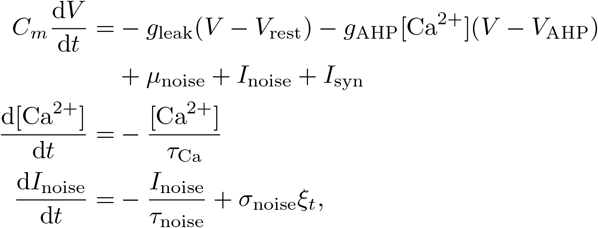

where *ξ*_*t*_ describes a white noise process. While Ren et al. [41] have modeled the spontaneous *I*_noise_ as pink noise, here we approximate its dynamics as an Ornstein-Uhlenbeck process, which represents another common model for neuronal input [94, Chapter 8]. This approximation allowed simulating different dynamical regimes more systematically by changing the mean input *µ*_noise_ and the noise amplitude *σ*_noise_ (see Fig. S5 **A**-**B**). The parameters describing the membrane potential dynamics were the membrane time constant *C*_*m*_ = 500 pF, the leak conductance 0.25 *µ*S, and the resting potential *V*_rest_ = *−*65 mV. For the after-hyperpolarizing (AHP) current, the conductance 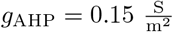, and the reversal potential *V*_AHP_ = *−*80 mV. The dynamics of the [Ca^2+^] were determined by the time-constant *τ*_Ca_ = 100 ms. When the membrane potential *V* surpassed the spiking threshold *V*_thresh_ = *−*50 mV, a spike was registered, and the potential was reset to *V*_rest_, and [Ca^2+^] was increased by 0.2 *µM*.

The synaptic input *I*_syn_ was composed of two sources: the intrinsic spiking activity of the network and the externally applied spiking activity. As an external spiking activity, we used spike-trains that were obtained from *in vitro* HD-MEA networks recording, which mimicked the studied experimental conditions. As in the study by Ren et al. [41], the synaptic effect of spikes at time 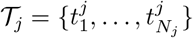 neuron *j* connected to neuron *i* was modelled as

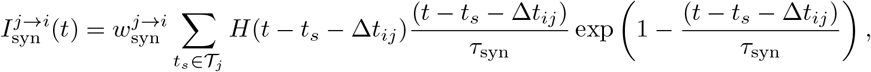

where *H*(*·*) was the Heaviside function. The total synaptic input to a neuron *i* was then given by the sum of all synaptic currents in Eq. S1A the neuron was connected to. The connection probability among simulated LIF neurons was set to 5% and HD-MEA input neurons were connected to LIF neurons with a probability of 10%. The synaptic weights were drawn randomly from a log-normal distribution 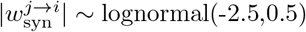, and the weight was positive when *j* was an excitatory neuron and negative for inhibitory neurons. If the synaptic weight exceeded the interval [0.05 nA, 0.4 nA], it was set to the limits of this interval. Δ*t*_*ij*_ was the synaptic time delay, which was given by distance_*ij*_*/*conductionvelocity. The distance between neurons distance_*ij*_ was the Euclidean distance between the positions of neuron *i* and *j*. The positions were drawn at random and uniformly from the rectangular area of the units from the HD-MEA input recording. The conduction velocity for each neuron was drawn uniformly from the interval 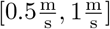. For the *in silico* benchmarking analysis across different inference algorithms (see Fig.2), we used 1 h-long simulations.

### S1B Simulations to validate connectivity inference from parallel HD-MEA/path-clamp recordings

The LIF framework described in the previous section was further adopted to also validate the methods applied to uncover synaptic connectivity from parallel HD-MEA/patch-clamp recordings (see Section S2 and Fig. S5). For a detailed description of the HD-MEA/patch-clamp setup and the performed voltage-clamp (VC) measurements, we refer the reader to Sec. S3B, S3C, and S3E. To simulate the patch-clamp recordings at a control voltage *V*_*c*_ = *−* 55mV, we constructed smaller subnetworks (LIF networks composed of n=10 neurons) with membrane currents of

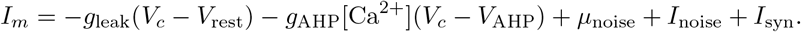

These neurons got input similar to all other neurons in the network, but since they were in VC-clamp mode, they could not spike. Next, we simulated 10 min of these combined data, i.e., registering spike times of the 300 LIF neurons as described in the previous paragraph and the currents of 10 VC-clamped neurons. The analytical results obtained from this analysis are shown Fig. S5 **C**-**D**.

## S2 Inferring synaptic connectivity from parallel HD-MEA/patch-clamp recordings

Next, we describe the modeling approach to derive synaptic connectivity statistically from parallel HD-MEA/patch-clamp recordings. Our model was inspired by previous work [75], with some notable modifications, in order to have only one statistical test per potential connection.

### S2A A regression analysis approach to estimate synaptic connectivity

In our model, we assumed that the recorded intracellular signal *y*_*t*_ is a linear superposition of two signals: i) a synaptic signal *y*^syn^ originating from synaptic signals from the extracellularly recorded presynaptic neurons, and ii), a residual signal *c*, that accounts for all intrinsic, and extrinsic fluctuations, such as synaptic signals from neurons that could not be sampled. Hence, the full signal was given by

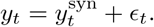

The modeled synaptic signal 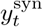 depends on the extracellularly recorded spiking activity of the presynaptic candidate neurons. Formally, the spike trains were a matrix with entries *s*_*t,i*_ = 1, if unit *i* elicited a spike at any time [*t*Δ, (*t* + 1)Δ); it was zero, if there was no spike. The synaptic signal *y*^syn^ was then modeled as a weighted sum of presynaptic signals ***x***

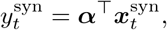

with coupling parameters ***α*** = (*α*_1_, …, *α*_*N*_)^*T*^. The presynaptic signals ***x***^syn^ were modeled the spike trains convolved with a response kernel *k*_*n*_(*τ*)

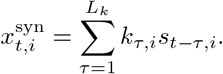

The response kernel *k* had the parametric form of an alpha function, which has been previously used for modeling the form of postsynaptic potentials (PSPs) [95, Chapter 5]. It was given by

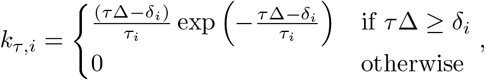

where *τ*_*i*_ the time constant, and *δ*_*i*_ the delay of the PSP. The latter is the main difference to Zhang et al. [75], which assumed a non-parametric footprint of the extracellularly recorded neurons on the intracellular signal. While the original model is likely more flexible, it requires multiple tests for each connection. In our model, however, we will subsequently only have one test per connection, i.e., whether the coupling parameter *α*_*n*_ ≠ 0. Everything, that could not be explained by the synaptic signal, such as transient fluctuations of the signal or the synaptic signals of unsampled neurons, we model by the following autoregressive process

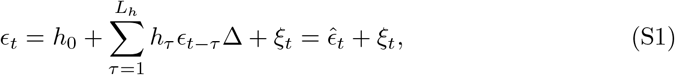

where *ξ*_*t*_ was a Gaussian noise with standard deviation *σ*_*y*_. Alternatively, *ϵ*_*t*_ can be written as 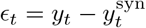. The former definition in Eq. (S1) only depends on the filter ***h*** and not on the couplings ***α***. The alternative definition of *c*_*t*_ is a function of ***α***, but not of ***h***. This fact allowed us to define the alternating optimization scheme described in the following section.

### S2B Fitting procedure

Given a recorded intracellular signal *y*_1:*T*_ and extracellular spike trains *s*_1:*T*,1:*N*_, we then sought to estimate the model parameters, i.e., the autoregressive filter ***h***, the synaptic coupling strengths ***α***, the kernel parameters ***τ*** = (*τ*_1_, …, *τ*_*N*_)^*T*^, ***δ*** = (*δ*_1_, …, *δ*_*N*_)^*T*^, and finally, the noise parameters *σ*_*y*_ by the maximum likelihood principle. The previously described model defined the following (log) likelihood for the observed data

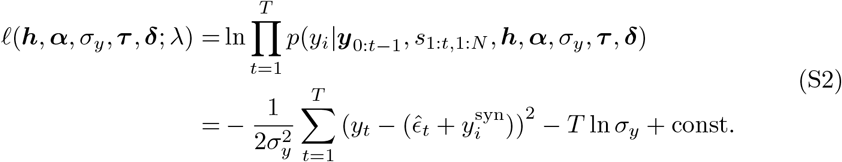

To avoid overfitting of the autoregressive filter ***h***, we included a regularizing term *𝓁*_reg_(***h***) to penalize non-smooth filters. Formally, the regularization term is the second derivative of ***h***, i.e.,

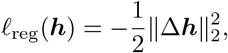

with Δ being the discrete Laplace operator. The optimal model parameters are given by

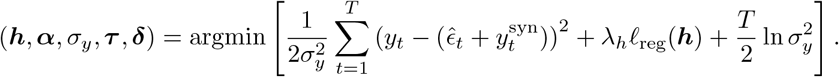

While there is no closed-form solution for this problem, we can derive analytic updates for the sub-problems by solving for ***h, α***, and *σ*_*y*_ separately. We, therefore, invoked an alternating optimization scheme. First, in order to find the optimal ***h***, we solved a linear problem. This problem is defined by computing the gradient of Eq. (S2) with respect to ***h*** and setting it equal to 0. In the same way, we then got the optimal couplings ***α***. *σ*_*y*_ can be similarly derived analytically. What remained was to find the parameters ***τ*** and ***δ***, which were computed by gradient ascent maximization of Eq. (S2)). We then alternated the optimization procedure until the likelihood converged.

Finally, we sought to determine, which extracellular neurons (respectively their recorded spikes, *s*_1:*T,i*_) were *de facto* connected to the neuron for which we had modeled the intracellular signal *y*. In other words, we asked which couplings were significantly non-zero (our null hypothesis was, that neurons are not connected, formally *H*_0_ : *α*_*i*_ = 0). To test for this hypothesis, we approximated the covariance of our estimate by the inverse Hessian matrix of the regularized log-likelihood (see Eq. (S2)) and used the absolute z-score *z*_*i*_ = |*α*_*i*_|*/σ*_*i*_ as test-statistics. For the latter, 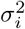 is the diagonal entry of the inverse Hessian matrix for couplings *α*_*i*_. For all couplings, where *z*_*i*_ ≥ *θ*_*α*_ the null hypothesis was rejected, i.e., these neurons were considered to be presynaptically connected to the patched neuron. The parameter *θ*_*α*_ was a threshold value that we derived by fitting the same model to jittered surrogate data (generated by adding Gaussian noise with a standard deviation of 5ms as jitter to the extracellular detected spike times). We took the 95% quantile of the z-scores *z*_*i*_, obtained by fitting the linear model to the jittered spike-train data, as a threshold.

## S3 HD-MEA recordings

### S3A High-density microelectrode arrays

To probe and record from *in vitro* developing primary cortical networks, we used complementary-metal-oxide-semiconductor (CMOS) based high-density microelectrode arrays (HD-MEAs). These chips comprise 26,400 platinum microelectrodes (size of electrode: 9.3 *×* 5.3 *µ*m^2^), with a 17.5 *µ*m pitch and a total sensing area of 3.85 *×* 2.10 mm^2^. HD-MEAs, as used in this study, allow for recordings from up to 1024 readout electrodes at the same time. The used custom HD-MEAs were bonded to printed circuit boards (PCBs), and a biocompatible epoxy (Epo-Tek 353ND, 35ND-T, Epoxy Technology Inc., USA) was used to encapsulate the bond wires and protect them from the medium. To decrease the impedance and to improve the signal-to-noise ratio (SNR), electrodes were coated with platinum black – deposited from a solution of hexachloroplatinic acid (7 mM, Sigma-Aldrich) and lead (2) acetate anhydrous (0.3 mM, Sigma-Aldrich) in distilled water, as described previously [96].

Two different types of HD-MEA systems, with comparable technical specifications, were used in this study: For the parallel HD-MEA/patch-clamp recordings, we used a custom single-well HD-MEA chip [97]. For the data and analysis presented in Fig. 5, we used a commercially available 6-well HD-MEA plate by MaxWell Biosystems (Zurich, Switzerland). The extracellular signals were acquired at a sampling rate of 20 kHz, for the single HD-MEA, and at 10 kHz, for the 6-well HD-MEA plate. Before the cell plating, we sterilized HD-MEA chips for at least 30 min in 70% ethanol and washed them 3 *×* with sterile deionized water; the electrode array was then treated with 0.05% (v/v) poly(ethyleneimine) (Sigma-Aldrich) in borate buffer (Thermo Fisher Scientific, Waltham, Massachusetts, United States) at 8.5 pH for 40 min and then washed 3 *×* with sterile deionized water.

### S3B Primary neuronal culture preparation

Rodent primary cortical neurons were prepared as previously described [96]: Cortices of embryonic day (E) 18/19 Wistar rats were dissociated in trypsin with 0.25 percent EDTA (Gibco), washed after 20 min of digestion in plating medium (see below), and finally gently triturated. Following cell counting with a hemocytometer, we seeded 15.000-20.000 cells (dataset 1: parallel HD-MEA/patch-clamp recordings ground-truth data; part of this data has been published in [58]), or 50.000 cells (dataset 2: HD-MEA network recordings; part of this data has been published in [98]) on each array, and placed it in a cell culture incubator for 30 min at 37^*°*^C/5% CO2. Then we added more plating medium carefully to each well (up until 1.5 mL). The plating medium was composed of: 450 mL Neurobasal (Invitrogen, Carlsbad, CA, United States), 50 mL horse serum (HyClone, Thermo Fisher Scientific), 1.25 mL Glutamax (Invitrogen), and 10 mL B-27 (Invitrogen). After two days, half of the plating medium was exchanged with maintenance medium. For the maintenance medium, we added 50 mL Horse Serum (HyClone), 1.25 mL Glutamax (Invitrogen), and 5 mL sodium pyruvate (Invitrogen) to 450 mL of D-MEM (Invitrogen). The maintenance medium was exchanged twice a week and at least one day before the recording sessions. All animal experiments were approved by the veterinary office of the Kanton Basel-Stadt and carried out according to Swiss federal laws on animal welfare. For dataset 1 (parallel HD-MEA/patch-clamp recordings), the experiments were performed on days in vitro (DIV) DIV16-18; for dataset 2 (1h-long HD-MEA network recordings), data were recorded at DIV14.

### S3C High-density microelectrode array recordings

In order to select active recording sites on the HD-MEA for long-term network recordings, we first recorded the multi-unit activity for each electrode across the whole chip using a series of dense-block configurations. Activity during this pre-processing step (’activity scan’) was assessed with an online sliding window threshold-crossing spike detection algorithm. The details for selecting the final recording electrodes are provided in the work by Bartram et al. [58] (dataset 1) and Akarca et al. [98] (dataset 2). Briefly, selecting a suitable network configuration involved, a ranking of the online detected mean spike amplitudes (per channel) and that channels showed a minimum of spike activity. Each HD-MEA network configuration consisted of approx. 1024 electrodes. The baseline recording for dataset 1 was composed of multiple network recordings on the day before the parallel HD-MEA/patch-clamp experiment (see Sec. S3D), yielding long network recordings of *>* 3 h duration. The network recording for the replication dataset (dataset 2) consisted of 1 h-long HD-MEA network recordings (n=6 cultures); here the network configuration was composed of up to 90 high-density electrode blocks (each block contained 4×4 electrodes).

### S3D Spike-sorting of HD-MEA network recordings

HD-MEA network recordings were spike-sorted using a semi-automated processing pipeline. For dataset 1 (HD-MEA/patch-clamp recordings), we combined the baseline recordings with the data obtained during the patch-clamp session. For dataset 2, we used the 1 h-long network recording. To spike sort HD-MEA network recordings, we applied the publicly available software package Kilosort 2 (KS2) [99], using parameters adapted to our data. Following spike-sorting with KS2, we manually reviewed all neuronal units deemed ‘good’ using the general user interface (GUI) of phy2 (https://github.com/cortex-lab/phy). We excluded units that showed aberrant spike waveforms, and that did not meet some standard quality criteria (e.g., less than 5% refractory period violations), or that had too few spikes (less than 1000 spikes). From the accepted ‘good’ units, we inferred whole-array spike-triggered electrical footprints and calculated all-to-all cross- and Pearson’s correlations to exclude potential duplicates. Split units, i.e., neuronal units for which the algorithm found more than one template, were also excluded from the data.

### S3E Whole cell patch-clamp electrophysiology

The parallel HD-MEA/patch-clamp experiments were performed using methodology introduced previously [57, 58]. To record from single neurons on the HD-MEA, we transferred the chips to a custom patch-clamp rig and perfused them with warmed (32-34 ^*°*^C) BrainPhys (BP). Patch-clamp recordings from cells located on the HD-MEA were obtained with borosilicate glass micropipettes (4-5 MΩ, Sutter Instruments, USA) containing (in mM): 85 caesium-gluconate, 60 CsCl, 10 Hepes, 4 Na_2_ATP, 0.3 GTP, 2 MgCl_2_, 0.1 EGTA, (pH 7.2-7.3; 280–290 mOsmol/l). Brief current-clamp recordings of spontaneous spiking were obtained, while synaptic activity was measured in voltage clamp mode at -70 mV holding potential. The high-chloride internal solution caused a shift in the GABA-A receptor reversal potential, which allowed us to record the synaptic activity of both GABAergic and glutamatergic synapses in one single recording. During the patch-clamp experiment, the same HD-MEA network configuration, as for the baseline recording, was used. This allowed to localize the patched cell on the HD-MEA, and to relate the intracellular obtained signals to the spike activity of the network. Patch-clamp recordings were carried out using an Axon Multiclamp 700B amplifier (Molecular Devices, USA), with digitization performed using an Axon Digidata 1440A (Axon Instruments). The recorded signals were low-pass filtered at 5 kHz and acquired with at least 20 kHz. Alexa 594 (20*µ*M, Thermo Fisher Scientific) was added to the internal solution to allow an assessment of the cell morphology. For details on the data, please see Bartram et al. [58].

## S4 Topological characterization of networks

Here we provide more details on the procedures on how we inferred connectivity from experimental HD-MEA derived spike trains, and how we analyzed this data using graph theoretical metrics (for details on the connectivity inference algorithms see *Methods and Materials* Sec. S5, and Fig. 2&5 in the main manuscript).

To assure reliable connectivity inference from the spike-sorted HD-MEA network data, we applied several filtering steps. First, we restricted our analysis to randomly selected units (100) that had at least 1000 spikes over the course of the recording. Second, we only estimated connectivity for edges that had a spike-sorting index *>* 0.5, as suggested by Ren et al. [41], and that had at least 200 spikes in the pairwise CCG of units – in a time window relevant for fast synaptic interactions (+/-20 ms). To generate jittered spike trains for units that passed the previously outlined quality criteria, and to be compatible with jittering code provided by [100], we removed any spikes occurencing at the zero-lag of the unit’s autocorrelation. Again, the aim of applying these arguably strict thresholds was to make sure that connectivity was estimated on a sufficient amount of activity and to reduce the likelihood of false positive connections.

Next, we selected several common topological metrics and compared them across network inference methods (see results presented in Fig. 2 and Fig. 5). We categorize these metrics broadly into global and local topological features. The global features describe average statistics for the given networks, while the local metrics describe topological values resolved per individual node. The global metrics included the overall *network density*, the network *global efficiency*, the average *clustering coefficient*, the *modularity* [101] and the *small-world index* [15] of the network and the occurrence of *triplet-motifs* in the data [77]. All topological features were calculated using algorithms provided by the `Brain connectivity toolbox` [102]. We briefly explain each metric below:

*Degree*. As the obtained binary graphs of the inferred networks were directed, the degree (*k*) denotes the sum of in- and outgoing edges of the observed network.

*Network density*. The network density was defined as the number of significant edges divided by the number of all possible edges in the respective network.

*Global efficiency*. The global efficiency (E) is calculated as the average of the inverse shortest path length (L). It is a measure of the global integration of a graph [103].

*Clustering coefficient*. The clustering coefficient (C) measures the clustering of connections/nodes in the network. C for directed networks is calculated as the fraction of realized directed triangles around a node, i.e., the observed number of triangles divided by the number of all possible triangles [104].

*Betweenness centrality*. The betweenness centrality (B) was defined as the fraction of all shortest paths in the network that contain a given node. Nodes with high values of betweenness centrality therefore participate in a large number of shortest paths [105].

*Modularity index*. The modularity index, Q, indicates how well a network can be partitioned into subgroups. We calculated it as proposed by [106].

*Small-word index*. The small-world index (S) of a binary network is usually defined by estimating two parameters: the characteristic path length of the network (L) and its average clustering coefficient (C). Both measures are normalized to appropriately randomized surrogate networks with the same number of nodes and edges [107]. Since some inferred networks contained disconnected nodes at the applied adoptive thresholds, and S is defined for connected networks, we used a variant of the small-world index [103]. We calculated S by dividing the normalized clustering coefficient (*C*_norm._) by the inverse of the normalized global efficiency (1*/E*_norm._).

*Motifs*. The frequency of triplet motifs was analyzed as proposed in previous work [77]. As for the small-world index, we generated appropriately randomized surrogate networks with the same number of nodes - and compared the empirical motif statistics to these randomized values. We compared a total of 13 different motifs.

To compare local topological features between the inferred networks and the LIF ground truth networks (see Figure 2), we computed Pearson’s *ρ* between the set of values of the true network and the estimated network. For quantifying the difference between global connectivity features, we calculated the relative difference (*y*_est_ *− y*_true_)*/y*_true_, where *y*_est_, and *y*_true_ are the feature of the estimated and true network, respectively.

## S5 Connectivity inference methods

In the following, we describe the connectivity inference algorithms implemented in this study. The considered algorithms include cross-correlogram (CCG)-based methods, methods based on information theory, neuronal synchrony, and finally, generalized linear model point processes. CCG-based methods have been widely applied to estimate synaptic connectivity from parallel recorded spike trains [33, 108]. Essentially, a CCG is a histogram of spike-time differences between two neurons *i* and *j*. We used the normalized CCG [30], defined as

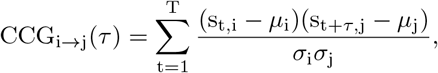

where *s*_*t,i*_ is the binary spike train of neuron *i* discretized in time bins with width Δ. *s*_*t,i*_ is 1 if neuron *i* spiked between *t* and *t* + Δ, and 0 otherwise. *T* is the number of time bins; *µ*_*i*_ and *σ*_*i*_ are the mean and standard deviation of the spike trains, respectively. To compute CCGs, we applied algorithms provided by the `Elephant` toolbox [109].

### Coincidence index

The first CCG-based method, implemented in this study, is termed *coincidence index* (CI, [56]). The CI was defined as

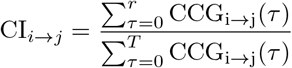

where *r* = *T*_syn_*/*Δ, and *T*_syn_ represents a time window in which synaptic effects are effective. Here we set Δ = 0.4ms and *T*_syn_ = 6ms. High CI values indicate an excess of spiking activity of neuron *j* after spikes of neuron *i*. As connectivity score, *CS*, we took the absolute z-score |CI_*i→j*_ − *µ*_*i→j*_|/*σ*_*i→j*_, where *µ*_*i→j*_ and *σ*_*i→j*_ are the mean and standard deviation of CI values obtained from surrogate spike trains of the corresponding neuron pair (50 iterations). The surrogate spike trains were generated by jittering the spike times of neuron *i* with uniform noise *U* (*−* 1.5*r*Δ, 1.5*r*Δ). The jittering strongly decreased the pairwise correlations observed for interactions in the synaptic time window. As putative synaptic weight, *W*, we simply took the value CI_*i→j*_.

### Smoothed CCG

While the CI relies on the generation of jittered surrogate spike-train data, Stark et al. [37] proposed a simple smoothing procedure in combination with a statistical test to assess whether the CCG deviates from the *H*_0_ hypothesis, i.e., that two neurons are synaptically not connected. Using this approach, the CCG is convolved with a Gaussian kernel, which is considered to have a similar effect on the CCG as obtaining a threshold value through spike-train jittering. Avoiding the jittering operation makes this approach computationally more efficient. We took the negative logarithm of the p-value as CS, and as weights *W* the synaptic strength, as described previously [36]. Throughout the manuscript, we referred to this algorithm as the *smoothed CGG* (sCCG) method.

### Generalized linear model CCG

An alternative method, combining the CCG approach with generalized linear models (GLMs), was proposed by Kobayashi et al. [40]. This approach decomposes CCGs into a slow and a fast fluctuating component. The model assumes that slow fluctuations, as observed in a pairwise CCG, can be regarded as the background activity within the network. Only fast short-latency fluctuations, that is, prominent peaks and troughs in the pairwise CCG that exceed the background activity, and that happen within the synaptic time window, should be considered as putative excitatory and inhibitory connections. Hence, Kobayashi et al. [40] proposed a parametric model, namely a GLM, to fit empirical CCGs. Formally, the GLM was given by

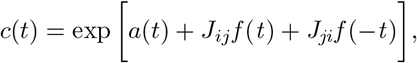

where *c*(*t*) is the co-firing rate for a timeshift bin *t. a*(*t*) models the slow fluctuation that encodes the background activity. *f* (*t*) models the synaptic interaction, which is given in the form of decaying exponential function 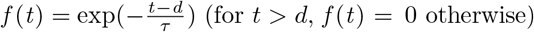, where *d* is the synaptic delay and *τ* is the time constant of the decay. The parameters *J*_*ij*_ and *J*_*ji*_ are the coupling strengths from neuron *i* to *j* and *j* to *i* respectively. Given a CCG from observed data, the maximum a posterior (MAP) estimate for parameters *θ* = {*J*_*ij*_, *J*_*ji*_, *a*(*t*)} is then obtained by numerical optimization. For details, the reader is referred to the original publication [40]. A similar approach has also been suggested by Ren et al. [41]. As *CS*, we take the z-score derived in [40, Eq. 14] given by 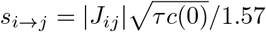. For the synaptic weight, Kobayashi et al. [40] proposed to transform the parameter *J*_*ij*_ heuristically to the size of a post-synaptic potential by the following formula *w*_*i→j*_ = *J*_*ij*_*/a*, where the factor *a* = 0.39, if there is a putative excitatory connection, i.e., *J*_*ij*_ *>* 0. On the other hand, for inhibitory connections, i.e., *J*_*ij*_ *<* 0, they used *a* = 1.57.

### Transfer entropy

Another important class of methods that has been widely applied to probe neuronal interactions, and to reconstruct neuronal networks, relies on information theory. The present study focussed on *transfer entropy* (TE) [54] and uses algorithms by the *IDTxl* toolbox [65] to estimate the functional connectivity between neurons. TE quantifies the “amount of predictive information” [110], between two processes – respectively, here, the spike trains of a source neuron *i* and a target neuron *j*. In brief, TE measures if including information on the spiking activity of neuron *i*, adds to the prediction of the future activity of neuron *j*, which goes beyond the information that is contained in the past activity of *j* alone. In the present study, we computed the TE on discretized spike train data with bins of size Δ = 5ms. As for the CI method, we used the absolute z-score *s*_*i→j*_ = |TE_*i→j*_ *− µ*_*i→j*_|*/σ*_*i→j*_ as CS. Again, *µ*_*i→j*_, *σ*_*i→j*_ are the mean and standard deviation of the TE values computed from jittered spike trains (50 iterations). The jitter noise was Uniform(*−*3.5Δ, 3.5Δ). As the weight of a connection, we considered the value TE_*i→j*_. Compared to the other implemented methods, the TE value is unsigned, i.e., it is always positive and, in that regard, could not distinguish between excitatory and inhibitory connections.

There are considerable limitations associated with the use of TE as a direct measure of coupling strength, as it may be confounded by the firing rate of neurons, the dynamical state of the network, the used embedding dimensions, and several other factors [110]. Future studies should implement and probe more recent TE variants that have been specifically developed for spike-train data [79] and that have addressed some of the limitations of current TE algorithms.

### Directed spike tiling coefficient

Cutts et al. [53] introduced the *spike time tiling coefficient* (STTC) to quantify synchronicity between spike trains. This method is computationally fast and has recently gained a lot of popularity. While the original method provided a measure of undirected pariwise correlation or functional connectivity, we modified the approach by Cutts et al. to a directed variant, which we call the *directed STTC* (dSTTC). In the following, we outline the dSTTC between the spike trains of neuron *i* and *j*. As in the original method, we defined a synaptic time window Δ_syn_ = 7ms. 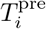 is the proportion of the total recording time, which is covered by time windows Δ before the spikes of neuron *i*. We note, that the times of overlapping windows are just considered once. Similarly, we define 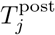 for the proportion of recording time, which is covered by windows Δ_syn_ following spikes of neuron *j*.

Furthermore, we define 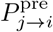 as the proportion of spikes of neuron *j*, that lies in the time windows Δ_syn_ preceding the spikes of neuron *i*. Similarly, 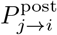 is the proportion of spikes of neuron *i* following the spikes of neuron *j*. Finally, we defined the dSTTC as

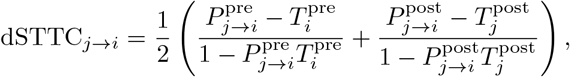

which can result in values in the range [ *−* 1, 1], as for the undirected original implementation by Cutts et al. [53]. The intuition behind the presented statistic is, that an excess of spiking of neuron *i*, that follows the spiking of neuron *j*, should indicate an excitatory connection. In this case, the dSTTC attains positive values. On the contrary, for inhibitory connections, we would expect a scarcity, or reduction, of spiking instead. In the latter case, the dSTTC would then result in more negative values. If the spikes of neurons *i* and *j* are occurring randomly, the dSTTC is expected to be close to 0. It should be noted, however, that these statements are based on the assumption, that the recorded data is stationary, i.e., there are no gross fluctuations in the firing rates. In the experiments at hand, however, this is rarely the case, due to transients in the firing rate and/or network burst dynamics. To mitigate such effects due to violations of the stationarity assumption, we resorted again to absolute z-score values for *CS* and calculate 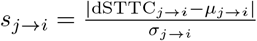, where *µ*_*j→i*_ and *σ*_*j→i*_ are the mean and standard deviation of the dSTTC values obtained from jittered spike trains. As in Cutts et al. [53], we used jitter noise Uniform(*−*3.5Δ_syn_, 3.5Δ_syn_). As weight *w*_*j→j*_ of a putative connection between two neurons, we took the raw dSTTC_*j→i*_ value.

### Point process generalized linear model

All presented algorithms so far were pairwise connectivity-inference methods. That is, they considered only two neurons at a time and neglected the potentially contributing effect of the activity of other neurons in their calculation. The *Generalized Linear Model Point Processes* (GLMPP) approach, however, is a framework that does consider such network interrelation – and has been previously used to probe connectivity [55]. This approach models the spiking of neuron *i* by a point process with rate *λ*_*i*_(*t* |*ℋ*_*t*_), where ℋ_*t*_ is the recorded spiking history up to time *t*. Here, we will assume that the rate model is

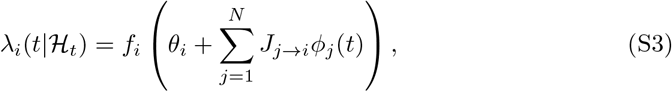

where the feature *φ*_*j*_(*t*) is the spike train of neuron *j* convolved with a causal exponential function with decay *τ* = 5ms. *f* is a monotonically increasing non-negative function. In this model, the parameters of interest are the coupling *J*_*j→i*_ for *i /*= *j*; note that*J*_*i→i*_ models how the neuron’s activity influences itself, such as, for example, the refractory period following a spike. For an excitatory connection *i → j*, we expected, that the rate *λ*_*i*_(*t*|*ℋ*_*t*_) increases after spikes of neurons *j*, and hence *J*_*j→i*_ should be positive. The contrary holds for inhibitory connections. If *J*_*j→i*_ was close to 0, this should indicate that no connection is present. We assumed a Gaussian prior distribution 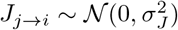 and 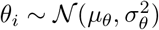. We intended to obtain the posterior distribution of the model parameters given the recorded spike trains. In general, this is not straightforward for a model defined by Eq. (S3). However, by choosing the *f* (*·*) to be a scaled sigmoid as in [72, 111], efficient variational algorithms have been developed to obtain an approximate Gaussian posterior distribution [73, 112] over the parameters *θ*_*i*_, and *J*_*j→i*_ via variational inference. Hence, once we have the approximate posterior density over parameters *θ, J*, we define the CS for a given connection as *s*_*j→i*_ = |*µ*_*j→i*_|*/σ*_*j→i*_, where *µ*_*j→i*_, *σ*_*j→i*_ are the mean and standard deviation of the posterior estimate for *J*_*j→i*_. As connection weight, we used the coupling value *J*_*j→i*_.

### Model specification and inference

Given the spike trains of several neurons, we can readily compute the features at any time

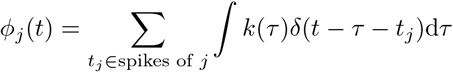

where 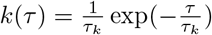 and *τ*_*k*_ = 5ms. In the following, we used the methodology of [73, 112] to fit the point process. In order to do so we needed to define the non-linearity *f*_*i*_ in rate in Eq. S3 as scaled sigmoid

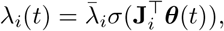

where 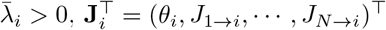 and ***θ***(*t*) = (1, *φ*_1_(*t*), *· · ·, φ*_*N*_ (*t*))^*T*^. For notational convenience, we dropped the conditioning on the history *ℋ*_*t*_. The likelihood of a point process [113] for spikes of neuron 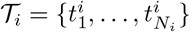 is

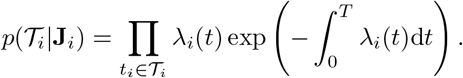

We assumed a Gaussian prior over the parameters **J**_*i*_ and a Gamma distribution prior over 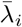. With this setting, we can utilize the augmentation scheme and the variational approach described in [112] to obtain an approximate Gaussian posterior over the **J**_*i*_, which we then used for the final connectivity.

## S6 Statistical thresholding of connectivity and evaluation of reconstruction performance

### Threshold selection

After inferring connectivity from either simulated or experimentally obtained spike-train data, several downstream analyses of this work required binary graphs. Hence, the connectivity matrices had to be thresholded. Selecting an appropriate threshold is a delicate task since it can affect the interpretation of the graph structure and its organizational properties. In the present study, we applied three different strategies. For the comparison of different inference methods on the LIF network data (see Fig. 2), we performed a search for the threshold that yielded the maximal Matthews correlation coefficient (MCC, see Sec. S6, i.e., the highest similarity to the underlying ground-truth graph. Such an approach has been previously applied in the literature [40], and does allow for a fair performance comparison across inference methods. However, since such a threshold optimization is not applicable to experimental data, we also report results using a second approach, that relied on a global adaptive thresholding logic (see Fig. 5 and S2). We, therefore, recalculated connectivity on jittered surrogate data (Gaussian jitter with a standard deviation of 10ms), and defined an absolute global threshold as the (1 *− α*) * 100% quantile of the resulting CS distribution. Here *α* can be interpreted as an expected false positive rate. The aim of this procedure was to destroy all short-latency synchronization by temporal jittering while keeping the firing rate dynamics intact. The jittered distribution of the CS values then reflected the null hypothesis, i.e., that there were no connections. For the HD-MEA recordings, we varied *α*-values from 0.05 to 0.001 (see Fig. 5). The data was only jittered once, which made this approach computationally fast. Finally, to show that topological results were stable across the selected statistical thresholds, we also applied proportional thresholds (see Fig. S4). With these thresholds, we probed graph metrics at a specific network density (e.g., 5%) and compared the topological properties of networks at a defined percentage of the strongest connections.

### Performance measures

To quantify the network-reconstruction performance across all inference algorithms, we applied standard validation measures, commonly used for classification tasks. The first metric is the *average precision score* (APS), which is threshold-free, i.e., there is no need to specify a threshold. The APS is calculated from the area under the *precision-recall* curve and is formally defined as APS = Σ_*n*_(*R*_*n*_ *− R*_*n−*1_)*P*_*n*_. *R*_*n*_, *P*_*n*_ are recall and precision if the *n*^th^ smallest CS would be selected as threshold. The APS provides values between 1 (perfect classification possible) and 0 (no connection is correctly classified without misclassifying all unconnected pairs). As a second performance measure, we implemented the *Matthews correlation coefficient* (MCC). The MCC requires a binarized connectivity matrix, respectively matrices, to compare networks. It is defined as

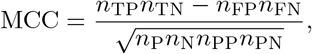

where *n*_TP_, *n*_TN_ are the number of true positives and false negatives, respectively, i.e., it is a measure of whether the algorithm classified the putative connections correctly. *n*_FP_, *n*_FN_ denote the numbers of false positives and false negatives. *n*_P_, *n*_N_ are the total number of connections and unconnected pairs of the ground-truth data. *n*_PP_, *n*_PN_ are the number of predicted positives (connection) and predicted negatives (non-connections), respectively. The MCC gives values between 1, for perfect classification, and *−*1 for the worst outcome.

**Fig S1.**
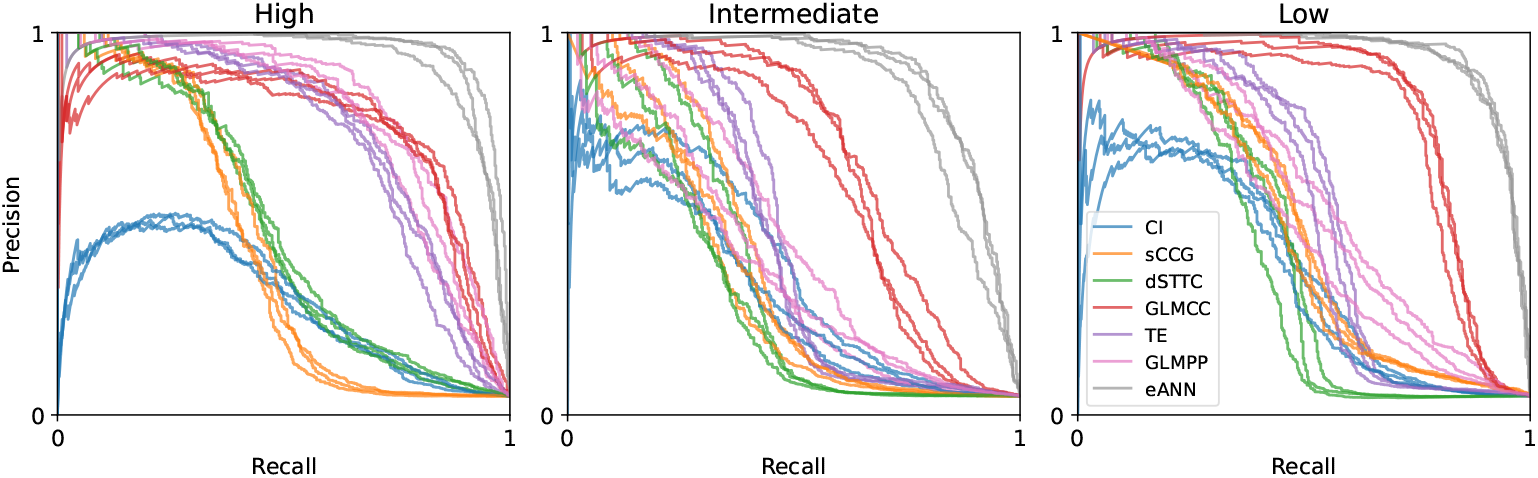
Precision-recall curves on simulated data. Curves for all methods on the data for Fig 2 (three lines per method for three networks (*N* = 100) that we fitted per High, Interm., and Low condition.). The more these curves extend to the upper right corner, the better classifications can be achieved. The eANN is the most robust method across all three conditions. The APS in Fig 2**D** is the area under these curves.

**Fig S2.**
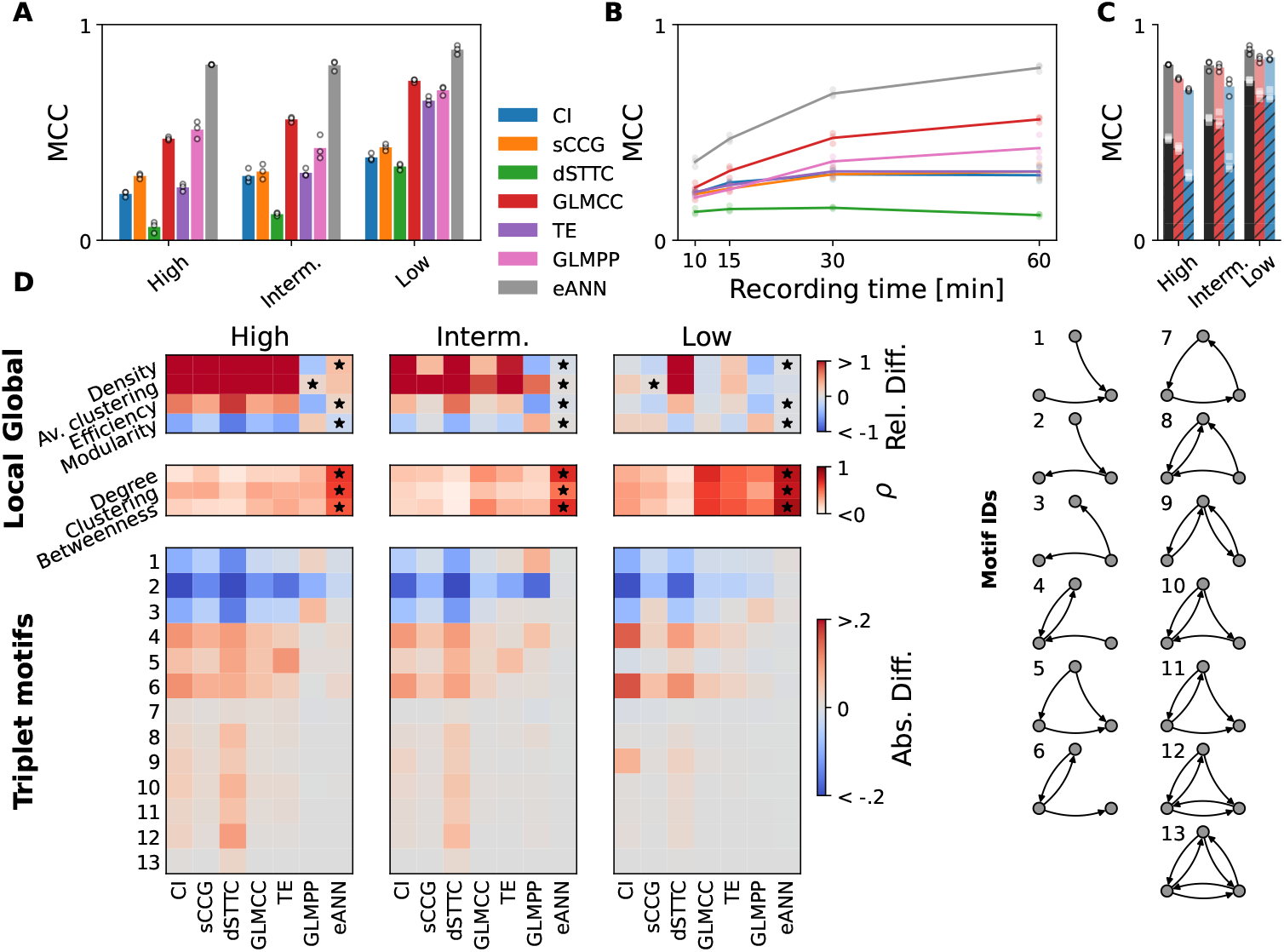
Network-reconstruction performance with global adaptive threshold. Results corresponding to Fig. 2 achieved with adaptive-threshold-selection (see Methods). **A** The MCC of different connectivity methods. Dots depict the performance obtained from fits on three different subnetworks of the same simulation. **B** Classification performance (MCC) as a function of recording time. **C** MCC for each type of connectivity, that is, excitatory (E, in red), inhibitory (I, in blue), combined (E+I, in black). Correspondingly, the performance gains achieved by the eANN are plotted in shades of red, blue, and black. **D** Quality of topological feature reconstruction for the inferred network across the three dynamical regimes. In the upper panel, the relative difference between four global features (network density, av. clustering, and efficiency) is shown. In the center panels, the Pearson correlation coefficient for local (per node of the network) features between the true and the inferred network is shown. Black stars indicate the method that performs best. In the lower panels, the absolute difference of triplet-motif frequencies between ground truth and the different estimated networks is shown.

**Fig S3.**
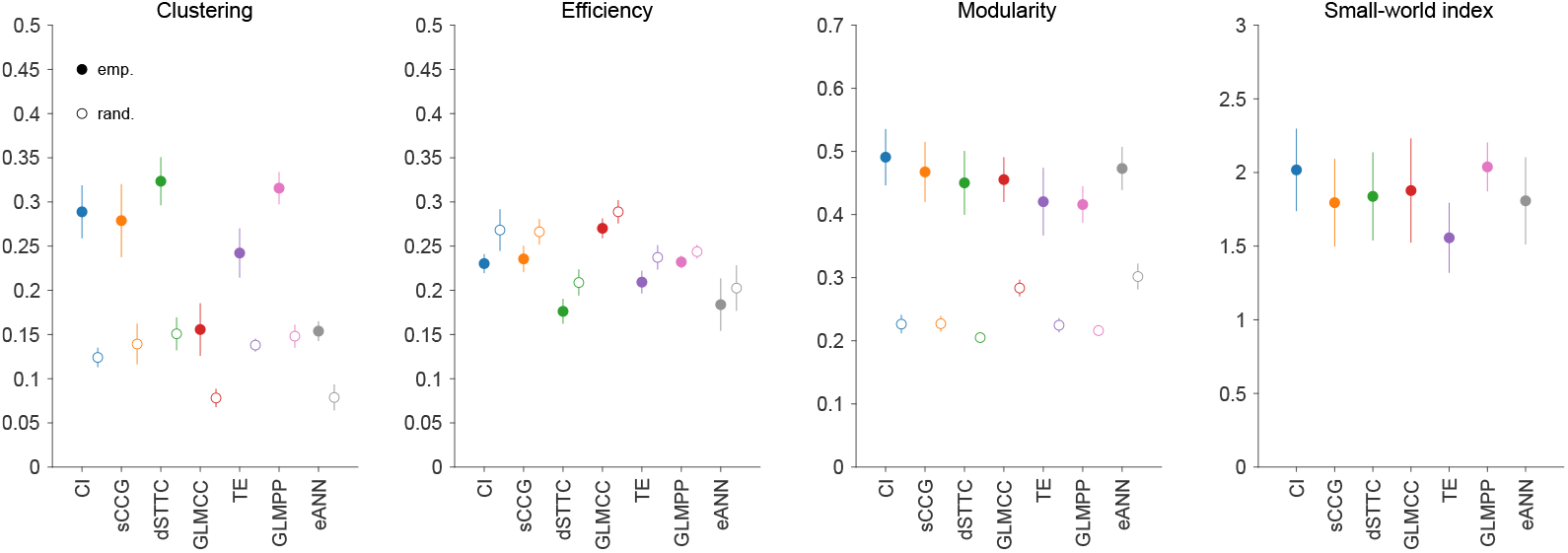
Comparison of *in vitro* neuronal network topology across inference methods. Four topological metrics (average clustering, efficiency, modularity, and small-world index) inferred from in vitro neuronal networks thresholded for the strongest 5% of connections in the network (proportional thresholding). Each panel depicts one topological measure; the colors correspond to the seven inference algorithms; colored circles correspond to the values obtained from the empirical data; the white-filled circles correspond to the surrogate networks (randomly rewired networks). In contrast to the analysis in Fig. 5**H**, this figure depicts all networks with the same number of edges.

**Fig S4.**
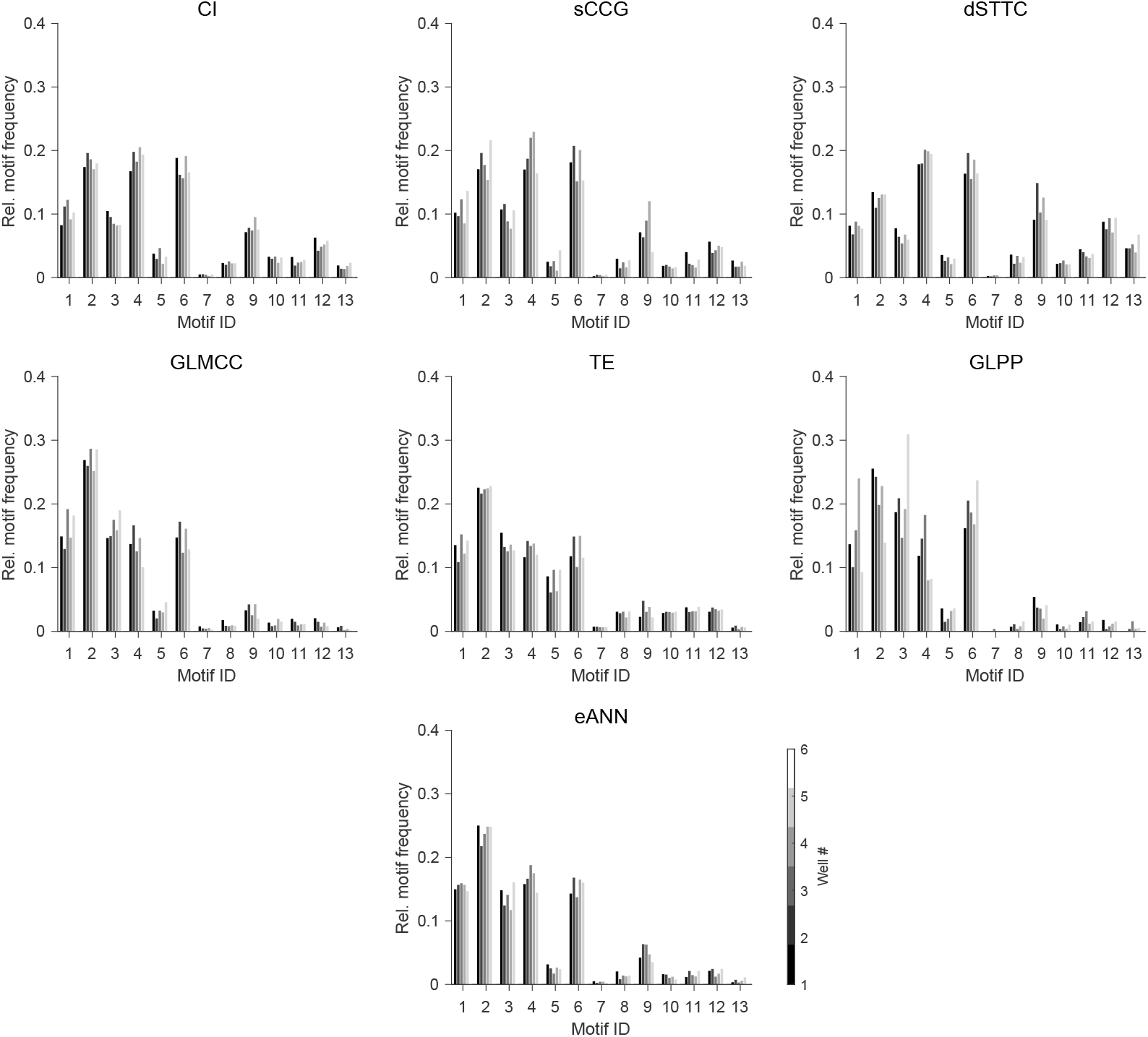
Triplet-motif frequency across inference methods and in vitro neuronal networks. The triple-motif frequency, observed in DIV14 *in vitro* developing neuronal networks (dataset 2), was very similar within each method class but differed considerably between some inference methods (see Fig 5**I** and **J**). Depicted motif frequencies were normalized, for each culture, by the total amount of observed motifs (*α* threshold: 0.01; network size: 100 units/network; 1 h recording duration).

**Fig S5.**
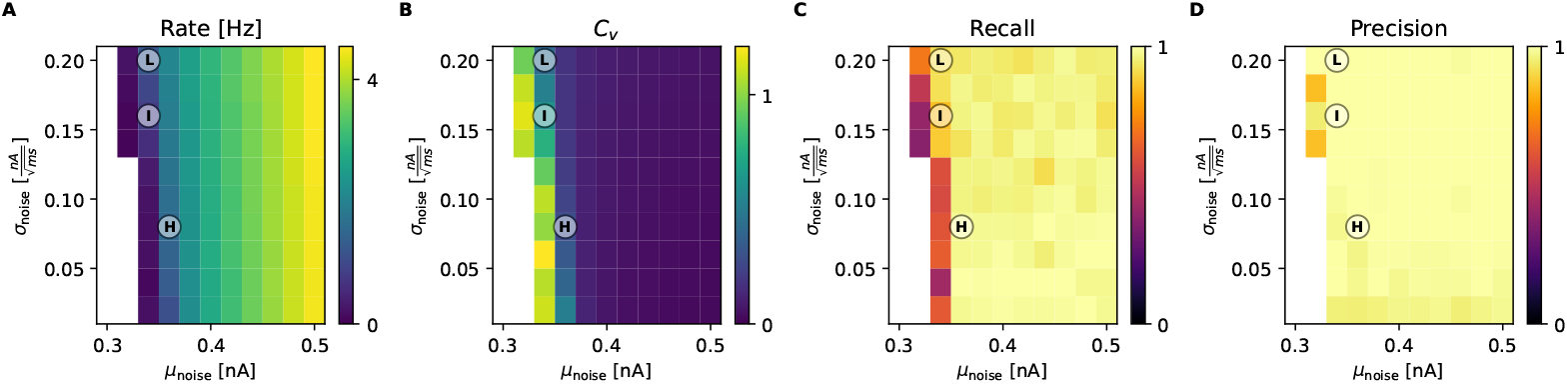
In silico validation of synaptic connectivity inference from simultaneous extra- and intracellular recordings. **A-B** An overview of the modeled dynamic range to validate the PSC connectivity inference method. The average spiking rate and Fano factor are shown for different simulations. Simulations with a rate *<* 0.5 Hz Hz were excluded. Circles correspond to simulations with **L**ow, **I**ntermediate, and **H**igh burst rate in Fig 2. **D-E** Display of recall and precision values for the IC connectivity inference method. Performance was averaged across VC recordings from 10 different neurons.

**Fig S6.**
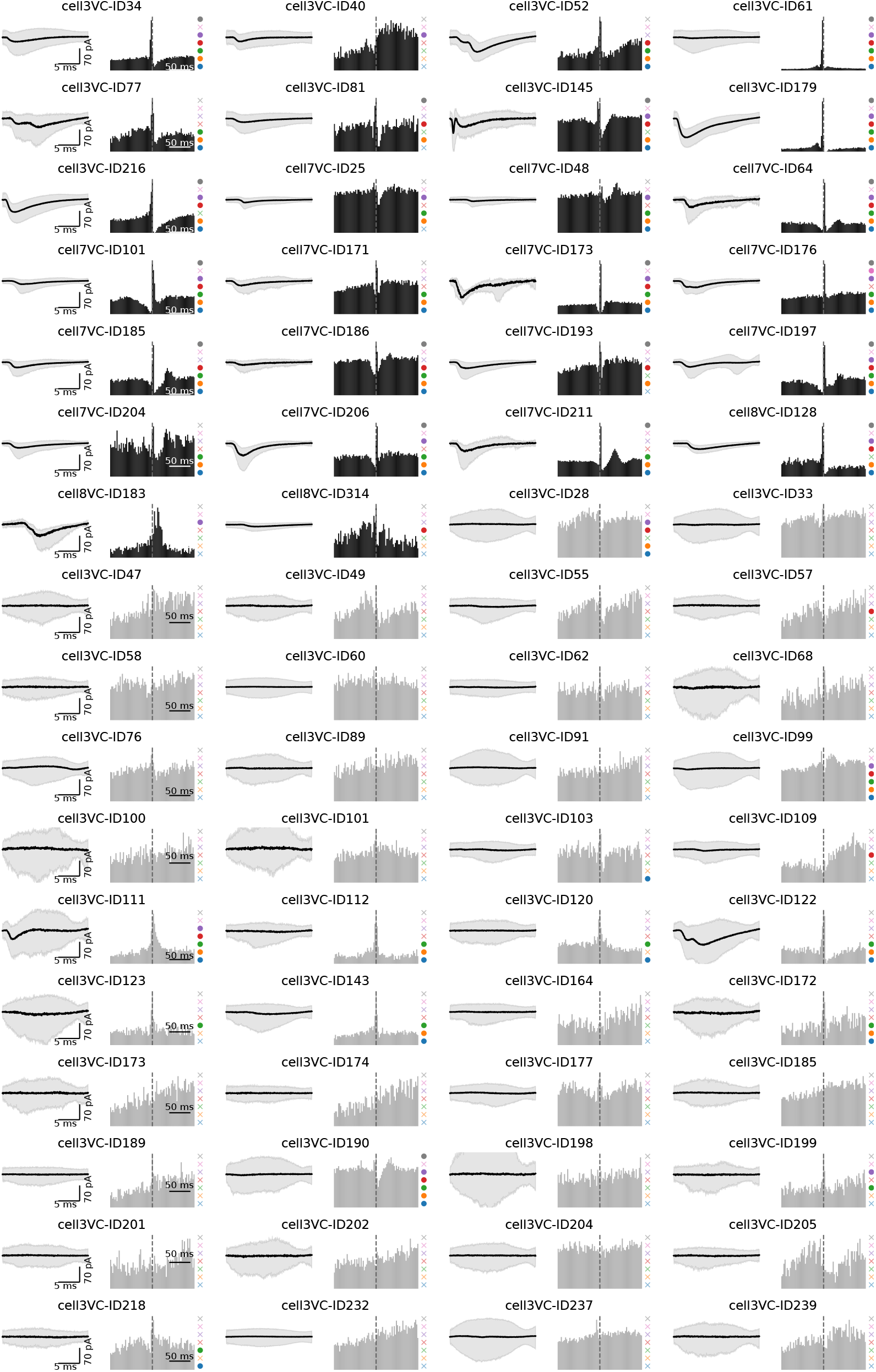
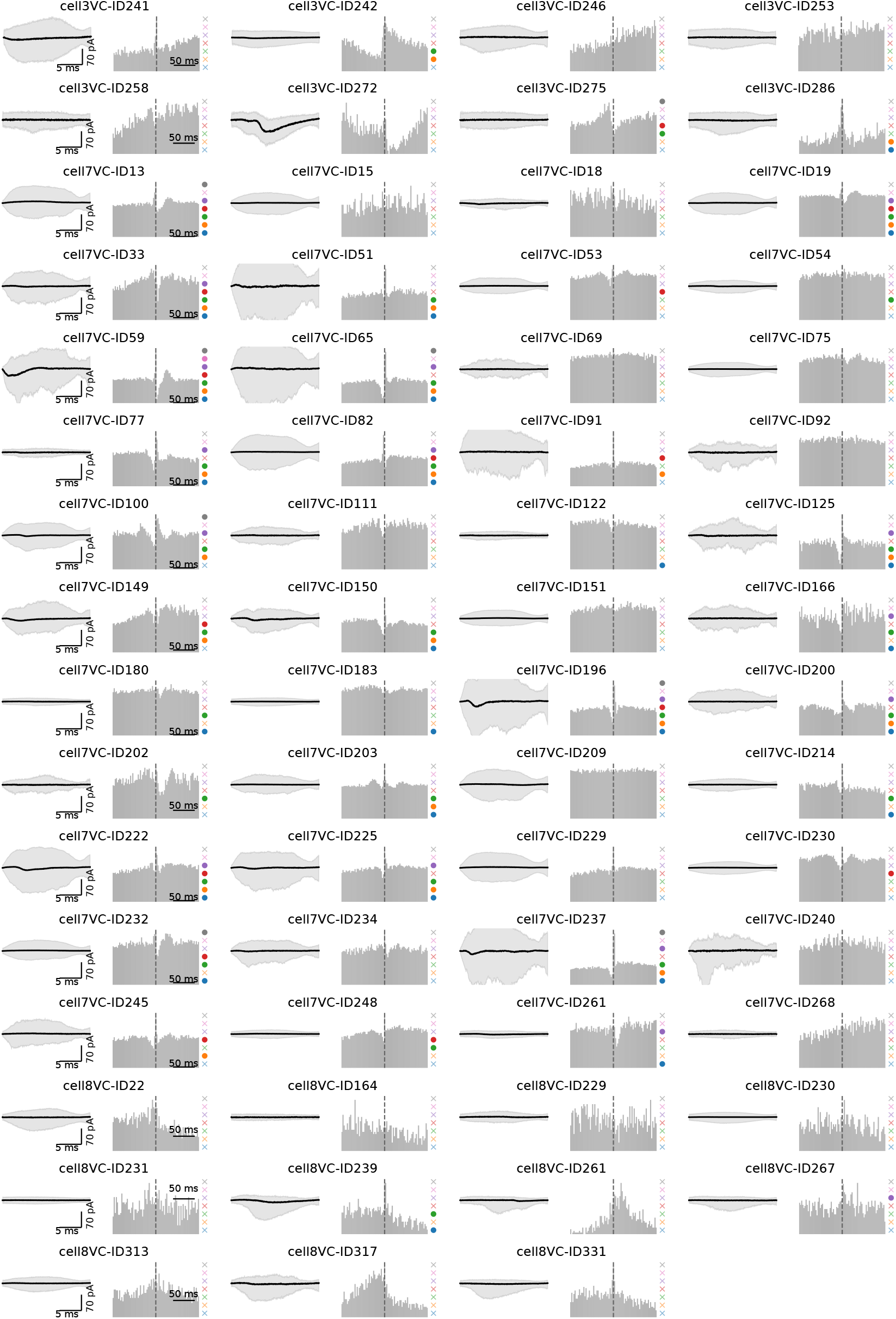
Connections in experimental ground-truth dataset. Here, we show all 26 experimental ground-truth connections found in the whole-cell patch recordings, together with all cell pairs, where no connection was found. We show the PSCs, the Spike-CCG, and the classification of the different connectivity methods similar to Fig. 4. Remaining cell pairs identified as non-connection.

**Fig S7.**
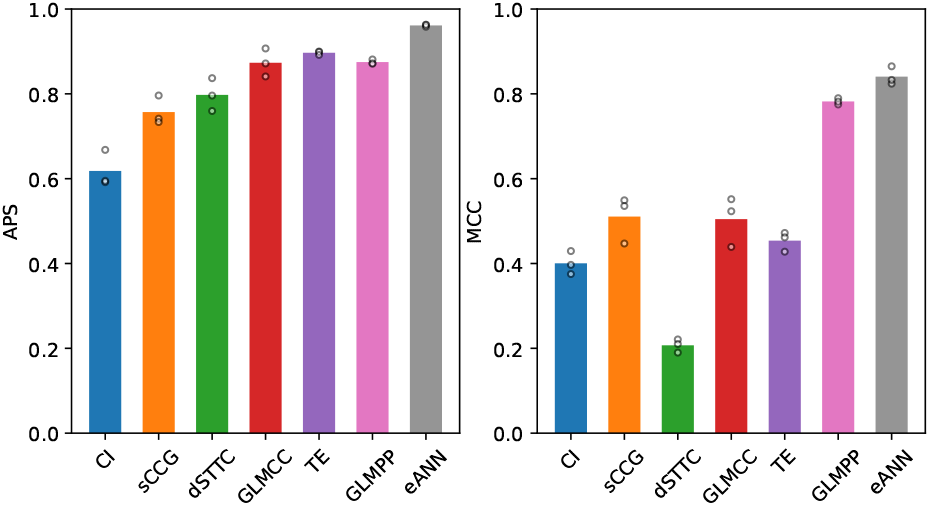
Performance on a dataset with different connectivity rule. Here, we show the results of the different connectivity inference methods on data simulated with 80/20 excitatory/inhibitory neurons. Population rate = 1.65Hz, Burst Rate=0.33Hz.

**Fig S8.**
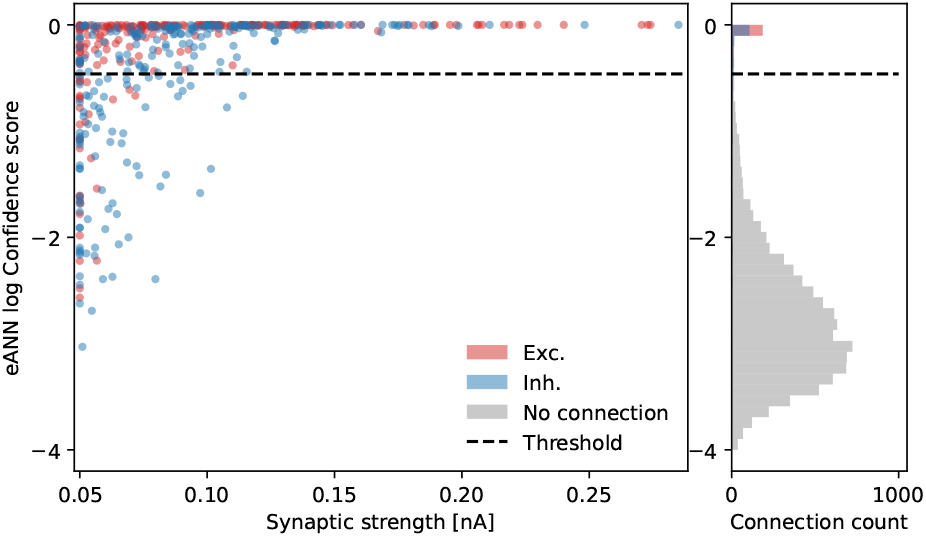
eANN confidence score. In the left panel the synaptic strength from the LIF simulations is plotted against the log confidence score of the eANN. We see lower confidence scores are more likely for weaker synaptic connections. Right panel shows a histogram of the log confidence score. This plot is done for data from Fig. 2 for the intermediate burst regime.

**Fig S9.**
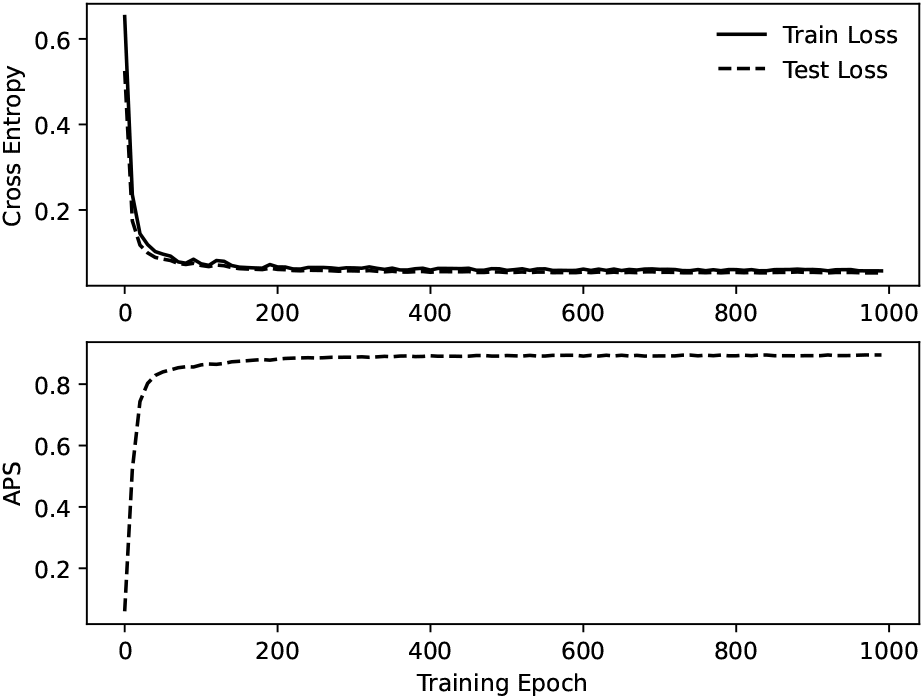
Training and test loss during eANN training. In the top panel, the cross-entropy loss is shown for the training set and a test set (intermediate burst regime). In the lower panel, the APS is shown on the test set during training.

**Fig S10.**
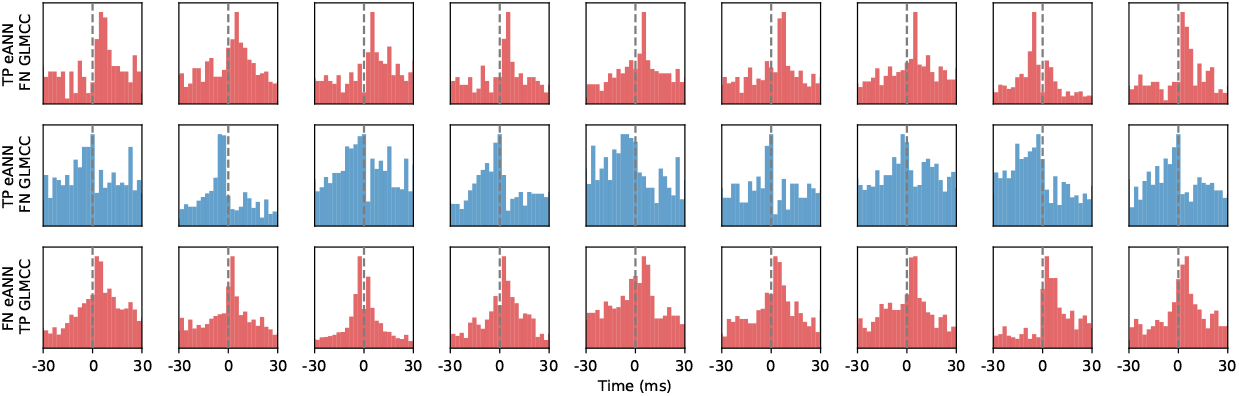
Comparison of eANN and GLMCC inference on synthetic data. Example cross-correlograms to compare connectivity inference of the eANN and the GLMCC method on some selected pairs (TP = true positive, NF = false negative). The bar color indicates if the ground truth connection was excitatory (red) or inhibitor (blue).

**Fig S11.**
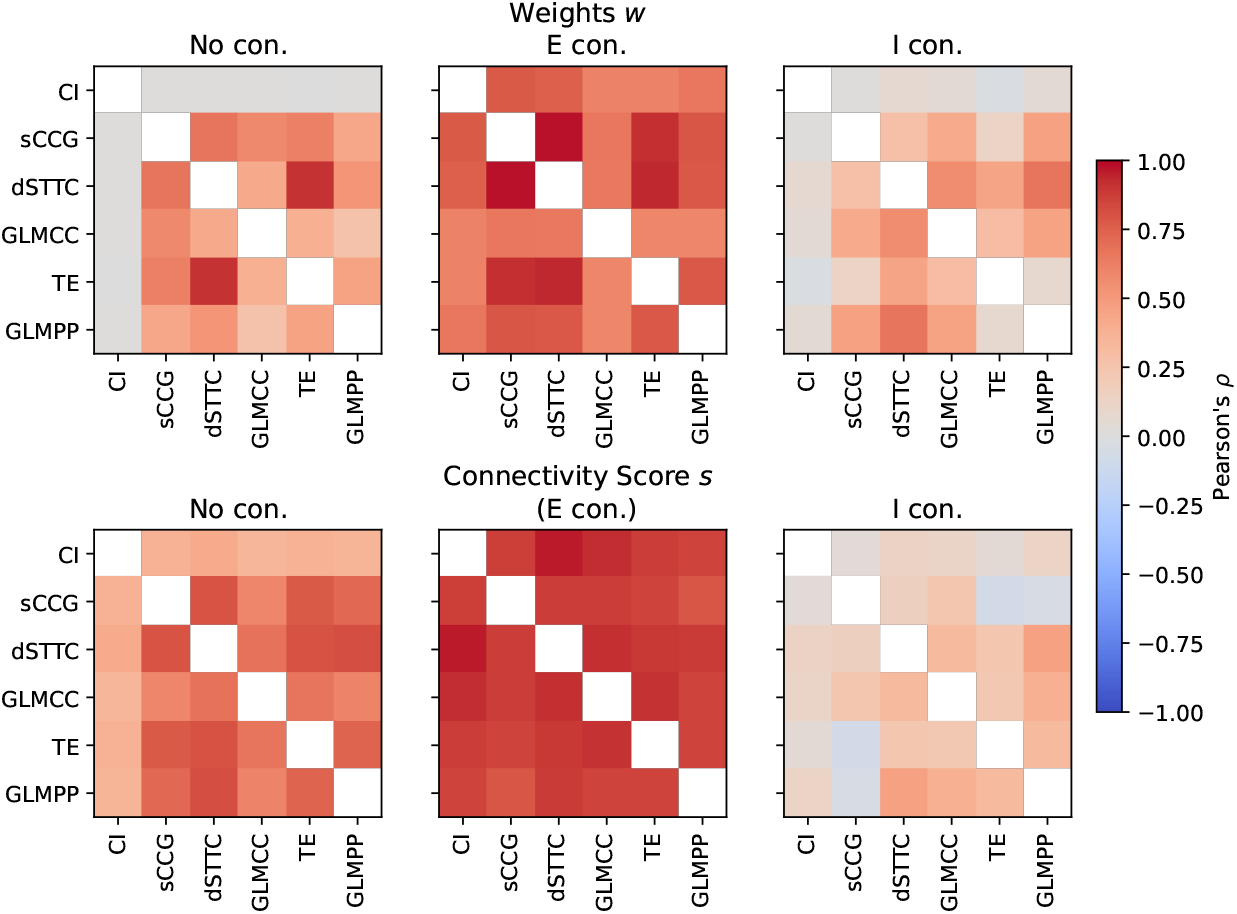
Pearson correlation of weights and connectivity score between different input methods. Left, middle, and right panel show the correlation for pairs of no, excitatory, and inhibitory connection, respectively. The data for the intermediate burst regime from Fig. 2 were used.

**Table S1.** Training simulations for the eANN.

| $\mu_{\text{noise}}$ [nA] | $\sigma_{\text{noise}}$ [$\frac{\text{nA}}{\sqrt{\text{ms}}}$] | Spike rate [$\frac{\text{spikes}}{\text{s} \cdot \text{neuron}}$] | $C_v$ | Burst rate [$\frac{\text{Bursts}}{\text{s}}$] |
| --- | --- | --- | --- | --- |
| 0.34 | 0.2 | 1.4 | 0.8 | 0.1 |
| 0.34 | 0.16 | 1.2 | 1.1 | 0.1 |
| 0.36 | 0.08 | 1.7 | 0.6 | 0.8 |
| 0.42 | 0.08 | 3.2 | 0.2 | 5.6 |
| 0.4 | 0.18 | 2.8 | 0.3 | 3.6 |

## Acknowledgments

This work was funded by a Swiss Data Science Center project grant (C18-10), by the two Cantons of Basel through a Personalized Medicine project (PMB-01-18) granted by ETH Zurich, and the European Research Council Advanced Grant 694829 neuroXscales’.

